# A selective sweep likely linked to xenobiotic detoxification identified along altitudinal gradients in the pine processionary moth

**DOI:** 10.1101/2025.11.24.690095

**Authors:** Pierre Nouhaud, Mathieu Gautier, Alexandre Schifano, Laure Sauné, Carole Kerdelhué, Charles Perrier

## Abstract

Understanding how natural populations adapt to complex environmental gradients is crucial for predicting evolutionary responses to global change. The pine processionary moth (*Thaumetopoea pityocampa*), a major forest pest expanding northward and upward in Europe, provides an ideal model to explore genomic adaptation along altitudinal and/or latitudinal gradients. We combined pooled and individual whole-genome resequencing of four pairs of low- and high-elevation populations from Spain, Italy and France (mainland and Corsica) to detect signatures of local adaptation. Population structure analyses revealed strong differentiation among distant populations but limited divergence within altitude pairs. Genome scans identified a few candidate regions under selection, and we notably uncovered a large (∼1 Mbp) region showing reduced nucleotide diversity, negative Tajima’s *D*, and fixed allele differences between high- and low-altitude populations, consistent with a recent selective sweep. This region includes a cluster of cytochrome P450 genes and the voltage-gated sodium channel gene *para*, both involved in detoxification and insecticide resistance. Although signatures of selection were observed at this locus in two distant population pairs, no definitive evidence of recent introgression was found between these distant populations, suggesting independent evolution. Altogether, our results do not show clear evidence for altitudinal adaptation in *T. pityocampa*, but suggest that xenobiotic exposure could exert a strong selective pressure locally. Our study highlights candidate genes and regions for further investigation of resistance evolution in an insect species in the wild.

**Significance statement:** Understanding how species adapt to their local environments is critical for predicting their survival under changing conditions. Here we show that populations of the pine processionary moth exhibit clear differences in genetic composition between high- and low-elevation habitats at a genomic region containing genes involved in detoxification and insecticide resistance. These findings suggest that local environmental pressures, such as xenobiotic exposure, can drive rapid evolutionary changes, highlighting the importance of considering local adaptation when managing pest species and predicting their responses to environmental change.

## Introduction

Climate change exerts a strong pressure on the vast majority of life forms, its swift pace increasing decline rate and extinction risk in natural populations (Thomas et al. 2004; Bellard et al. 2012; Scheffers et al. 2016). To persist, species must either shift their distributions latitudinally and/or altitudinally, acclimatize to novel stressful conditions through phenotypic plasticity, and/or undergo rapid adaptation (Parmesan and Yohe 2003; Chen et al. 2011; Inouye 2022). These strategies are not mutually exclusive, and natural populations may often combine different responses to survive, for instance through local adaptation at expanding range margins (Hill et al. 2011). Yet, climate is not the sole driver of change, as colonization of new habitats also exposes populations to additional selective pressures, many of which may also be anthropogenically induced (for a review in insects, see McCulloch and Waters 2023). Population genetics provides a powerful framework to map signatures of selection along the genome, offering important insights into how natural populations may respond to climate change (Bernatchez et al. 2024). Numerous methods have been developed to identify candidate adaptive loci from sequencing data (reviewed in, e.g., Bourgeois and Warren 2021). Divergence-based genome scans, for instance, have been widely used to detect loci whose genetic differentiation exceeds expectations under neutrality (e.g., Dudaniec et al. 2018). Alternatively, genotype–environment association (GEA) analyses assess statistical relationships between allele frequencies and environmental variables to identify the drivers of local adaptation (Forester et al. 2018). Because both divergence and GEA scans are prone to type-I errors (see, e.g., Lotterhos and Whitlock 2015), taking into account the demographic history of populations (Bonhomme et al. 2010; Günther and Coop 2013) and incorporating replicated population pairs (e.g., Pélissié et al. 2022) altogether provide a robust strategy to more confidently detect candidate adaptive loci showing signatures of selection.

The pine processionary moth *Thaumetopoea pityocampa* (Dennis & Schiffermüller; Lepidoptera: Notodontidae) is one of the most severe pests of pine trees in Europe and North Africa (Kanat et al. 2005). Late instar larvae also represent a significant public health concern due to their urticating hairs, which can cause serious dermatological, ocular, and respiratory lesions in humans and animals (Rivière 2011; Vasseur et al. 2022). The potential pine processionary moth distribution is mainly constrained by winter temperature (but see Rossi et al. 2025) as larvae are unable to feed and eventually die of starvation below ca. -15°C (Robinet et al. 2015). While the species has been expanding across Europe since the last glacial maximum (Kerdelhué et al. 2009), in recent decades, rising temperatures have increased the expansion pace of populations northward across Europe (Battisti et al. 2005), along with a retraction of its southern range (Tunisia, Bourougaaoui et al. 2021). Whereas the northern distribution limit remained near 48°N until the 1980s, it now extends beyond 49°N (Robinet et al. 2025), and species distribution models suggest that the species could expand to 50°N under future climate scenarios (Rossi et al. 2025). In addition to its northward expansion, the pine processionary moth has been recorded establishing at higher elevations, with altitudinal colonisation rates estimated at approximately 70 meters per decade on south-facing slopes (29 meters per decade on north-facing slopes, Battisti et al. 2005). Although the latitudinal and altitudinal range expansion of the pine processionary moth across Europe is well documented, with, e.g., earlier emergence recorded at higher elevation levels (Martin et al. 2022), whether populations locally adapt to ecological variations on altitudinal gradients remains an open question.

Here, we take advantage of the recent release of a chromosome-level assembly (Gautier et al. 2026) to study potential genomic footprints of local adaptation to altitudinal ecological variation. We implemented a replicated sampling design, collecting individuals from low- and high-elevation sites colonised post-glacial warming (i.e., prior to the last few decades, see Martin et al. 2021) across four distinct localities around the Mediterranean basin (Figure 1). To obtain genome-wide data for each of these eight populations, we used a combination of pooled sequencing (pool-seq) and individual sequencing (ind-seq, see Table 1 for detailed sample sizes). This hybrid, cost-effective approach was adopted to ensure precise allele frequency estimation for association scans (pool-seq, see Futschik and Schlötterer 2010), while allowing reconstructing effective population size histories and identifying putative signatures of structural variation (ind-seq). We first characterized the genetic structure of pine processionary moth populations. We then applied model-based approaches to identify candidate loci associated with local adaptation and with altitude, hypothesizing that genes involved in the circadian rhythm and in metabolism regulation could represent promising candidates. While we did not detect convergent signals of association to altitude across the four altitudinal gradients, we uncovered putative signatures of selection in a region containing a cluster of cytochrome P450 genes and the voltage-gated sodium channel gene *para*, both involved in detoxification and insecticide resistance. Altogether, our results indicate that multiple selective pressures are acting in natural populations and highlight the need for careful interpretation of adaptive signals in genomic data.

**Figure 1.**
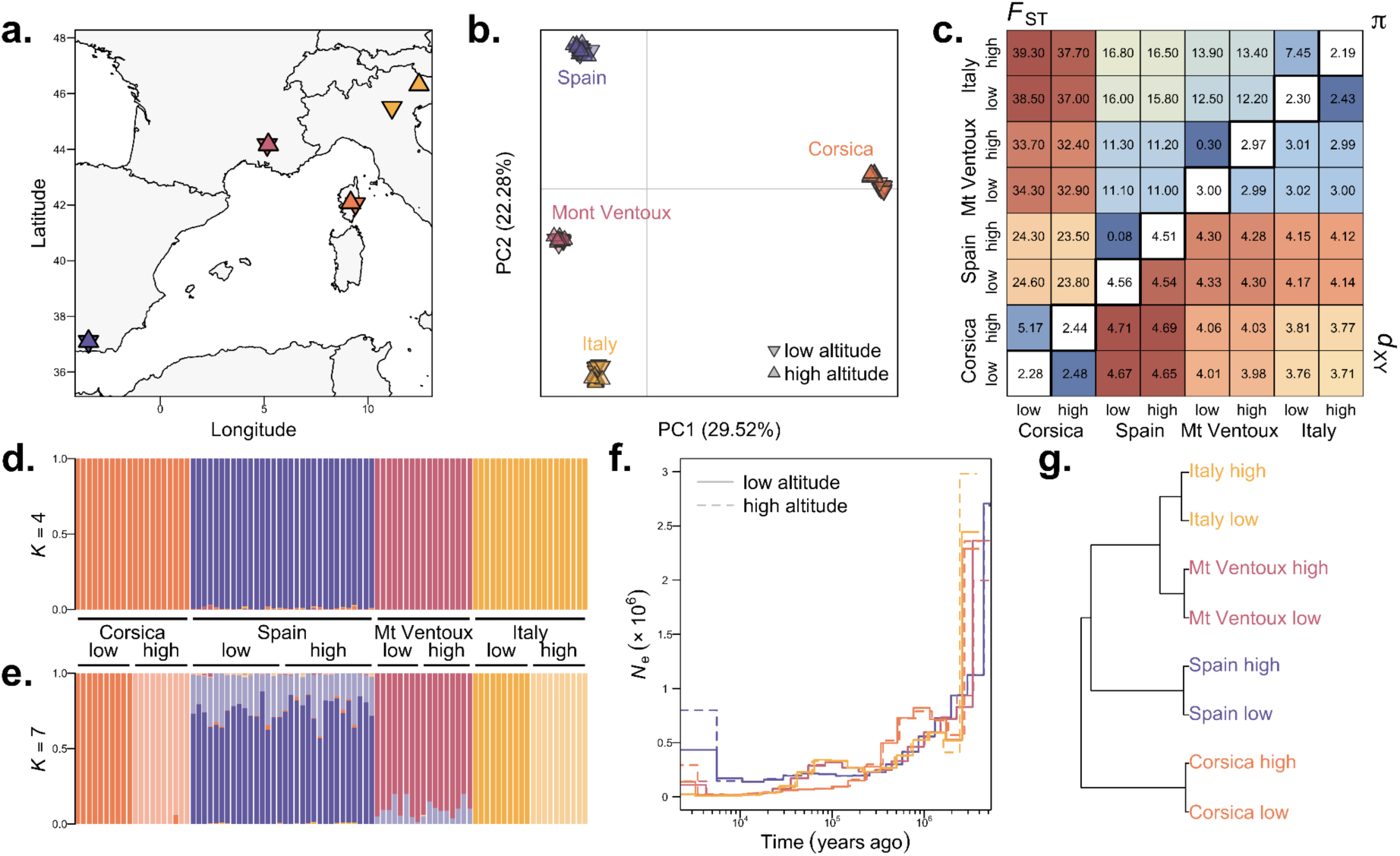
Sampling locations and genetic structure of the pine processionary moth individuals and pools. **a.** Map of the sampled localities. **b.** PCA of the 89 ind-seq libraries genotyped at 84,443 linkage-pruned autosomal SNPs. **c.** Genome-wide nucleotide diversity (π, ×10^3^, diagonal) and pairwise divergence metrics (*F*_ST_, ×10^2^, upper triangle; *d*_XY_, ×10^3^, lower triangle) computed on ind-seq libraries. **d.** and **e.** Individual sSupervised clustering results obtained with Admixture for *K* = 4 and *K* = 7 ancestry components, respectively (ind-seq data). **f.** Effective population size histories inferred from ind-seq data using MSMC2. **g.** Hierarchical clustering tree of the 8 pool-seq libraries based on the scaled covariance matrix of population allelic frequencies (Ω) estimated by BayPass.

**Table 1.**
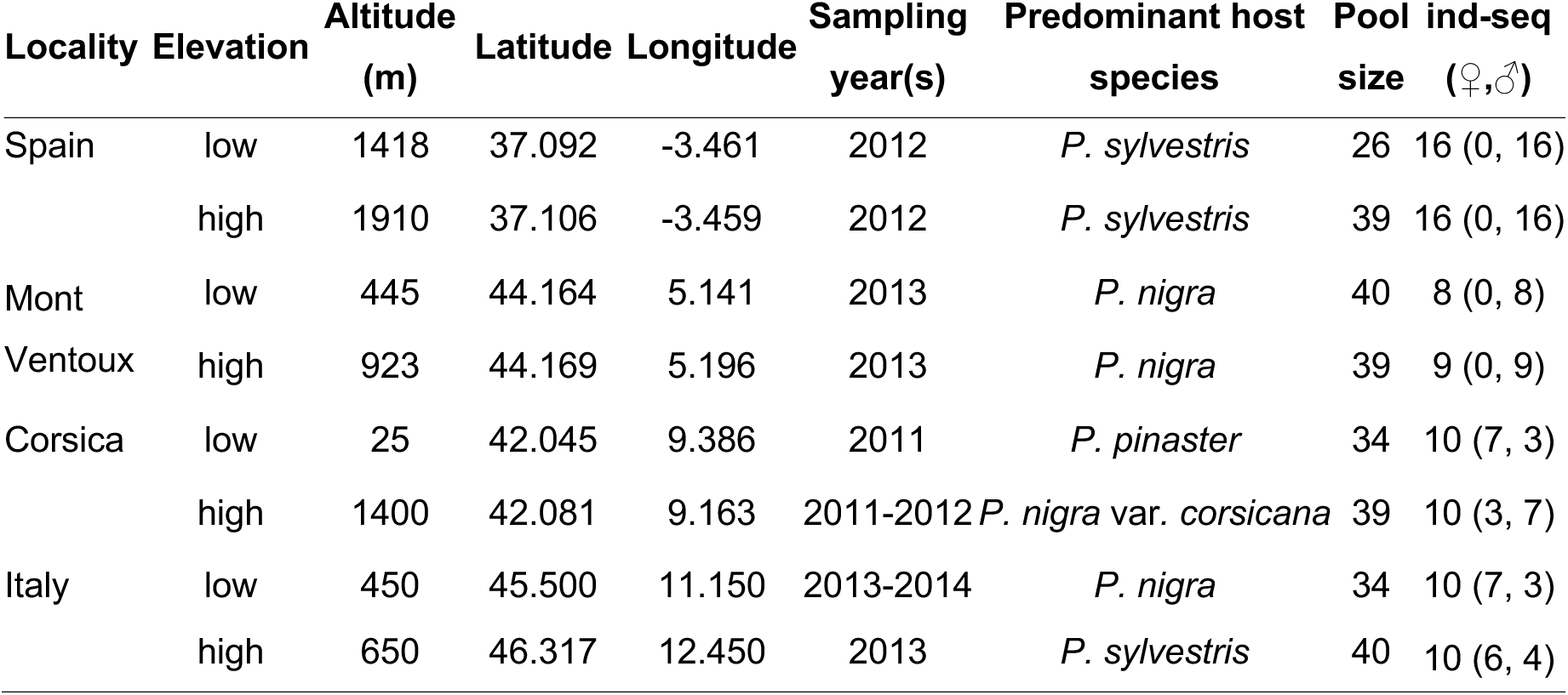
Overview of sampling locations, local predominant *Pinus* host species and associated data sets. Pool sizes are indicated as diploid, and numbers in brackets for ind-seq samples correspond to sex counts inferred from the Z-chromosome-to-autosomal depth ratio analysis (see Methods).

## Results

### Strong structure among distant populations but little differentiation within low and high altitude pairs

Sequencing produced between 92.3-682 (median: 123) and 260-577 (median: 374) million reads for 89 ind-seq and 8 pool-seq libraries, respectively, with mean insert sizes ranging from 340 to 457 (median: 387, see Suppl. Table 1 for detailed sequencing statistics). Mean depths ranged from 15.7× to 109× (median: 21.9×) for ind-seq libraries and from 41.4× to 97.7× (median: 60.4×) for pool-seq libraries. Heterosomal-to-autosomal depth ratios identified 66 males and 23 females from the ind-seq data (Table 1, Suppl. Table 1, Suppl. Fig. 1). Joint SNP calling and subsequent library-specific filtering recovered 14,323,239 sites for the pool-seq dataset and a subset of 9,193,257 sites for the ind-seq dataset.

The PCA carried on 84,443 linkage-pruned autosomal SNPs clustered samples by locality, with the first axis discriminating between Corsican samples and the rest (Fig. 1b). Nucleotide diversity measured with π was highest in the two Spanish populations, intermediate in Mont Ventoux and comparably smaller in Corsica and Italy (Fig. 1c). Overall, absolute and relative pairwise divergence were low to moderate between populations within localities, and the increased differentiation observed for the Italian pair could be caused by the greater distance between sampling sites (ca. 135 km, Fig. 1a). Divergence was however higher between localities, with *F*_ST_ values ranging from 0.11 (Mont Ventoux *vs*. Spain) to 0.38 (Corsica *vs*. Italy). Supervised clustering grouped individuals per locality at *K* = 4 (Fig. 1d), while Puechmaille’s method gave K = 7 as the most likely number of ancestry components (Fig. 1e). Under this scenario, both Corsican and Italian samples were split according to elevation levels, while Mont Ventoux samples remained within a single cluster and Spanish samples appeared as admixed, which could be a consequence of the strong nucleotide diversity present in these samples compared to the rest (Fig. 1c). The reconstruction of demographic trajectories through time confirms a relatively recent divergence between low- and high-altitude populations within localities, compared to between localities (Fig. 1f). Divergence between the ancestral Corsican population and mainland populations would have occurred during the mid-Pleistocene transition 0.8-1.2 million years ago, after which the ancestral Corsican population showed a declining effective population size. The separation of estimates between the ancestral Spanish population and both ancestral Italian and Mont Ventoux populations began about 250,000 years ago, while the split between these last two populations happened more recently, in line with their lower absolute divergence estimates (Fig. 1c). The scaled covariance matrix of population allelic frequencies Ω estimated from pool-seq data using BayPass recapitulated the population structure identified from ind-seq data, grouping populations per locality and suggesting higher divergence between the Corsican pair and the rest of the samples (Fig. 1g).

### Genomic footprints of divergent selection and of association with altitude

To identify footprints of divergent selection, including patterns of association with altitude, three distinct BayPass analyses were carried from pool-seq data while taking into account population structure (Fig. 1g) using 14.3 million SNPs genome-wide. For each analysis, local scores were computed to identify candidate regions genome-wide by combining SNP-specific statistics at 4,635,467 SNPs fulfilling a BayPass-estimated minor overall allele frequency above 10% (Fig. 2).

**Figure 2.**
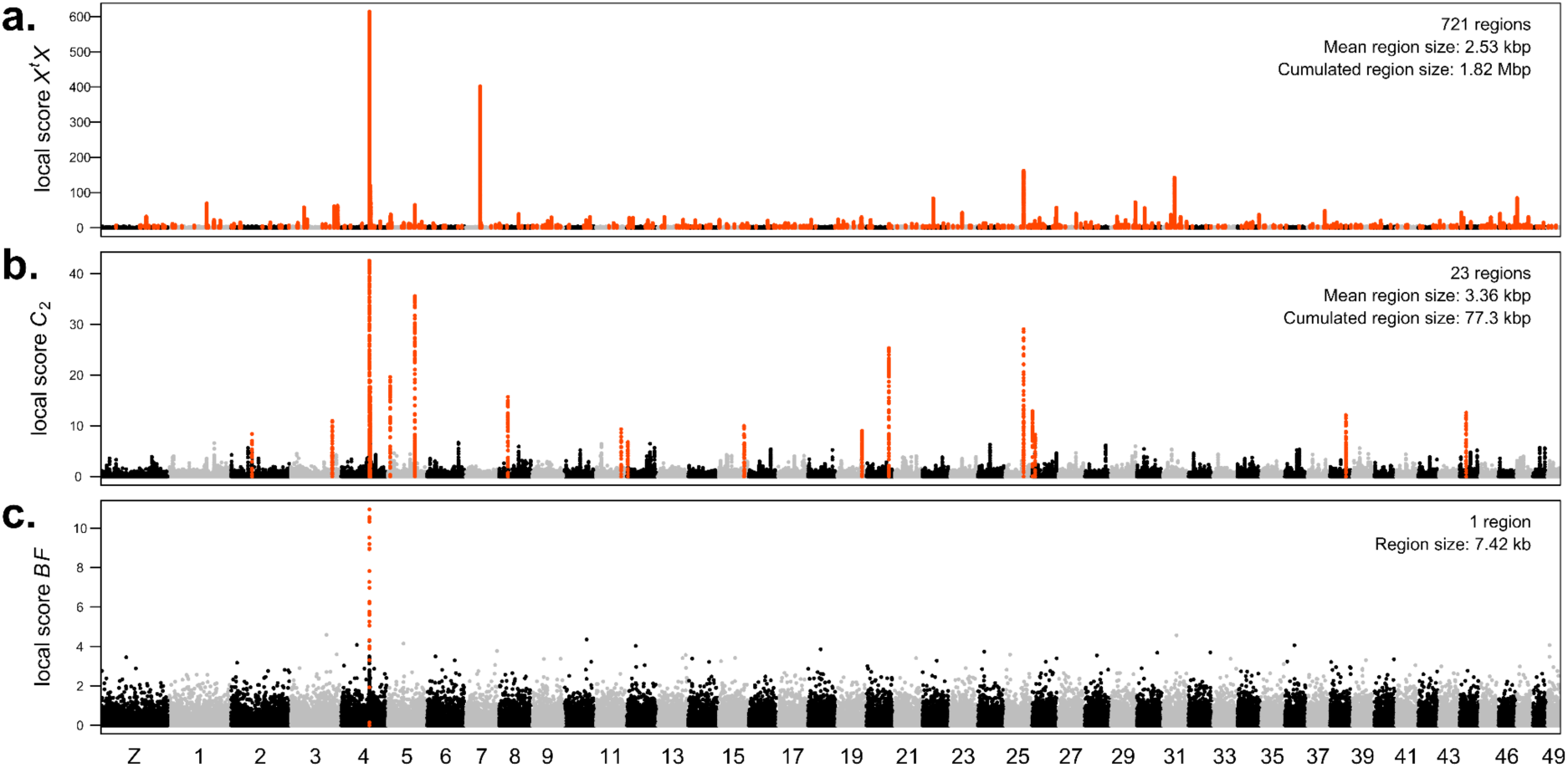
Results of genome-wide scans for signatures of selection at 4,635,467 SNPs (MAF ≥ 10%) genotyped in pool-seq libraries along the 50 chromosomes of the pine processionary genome. For each panel, SNPs located within significant local score regions are depicted in red, with region number and sizes provided in the upper right corner. **a.** Local scores of the *X^t^X* statistic detecting positive selection signatures, irrespective of the elevation levels (ξ = 3, significance level *α* = 0.001). **b.** Local scores of the *C*_2_ statistic, contrasting low and high altitude groups of populations (ξ = 2, *α* = 0.01). **c.** Local scores of Bayes Factors (*BF*) measuring the strength of evidence for the linear correlation between population allele frequencies and altitude (ξ = 2, *α* = 0.01).

The first analysis focusing on signatures of positive selection relied on the *X^t^X* statistic, a site-specific differentiation metric explicitly corrected for the scaled covariance of population allele frequencies (Günther and Coop 2013). Our *X^t^X* scan identified 721 genomic regions scattered across all 50 chromosomes, with a mean region size of 2.53 kbp (min: 23 bp, max: 134 kbp) and a cumulated size of 1.82 Mbp (ca. 3‰ of the genome, Fig. 2a, Suppl. Table 2). This set of regions overlapped with 227 unique gene models (Suppl. Table 3). The second analysis contrasted allele frequencies corrected for population structure between high- and low-elevation population groupings (*C*_2_ statistic, Olazcuaga et al. 2020) and identified 23 genomic regions scattered across 14 chromosomes, with a mean region size of 3.36 kbp (min: 420 bp, max: 11.4 kbp) and a cumulated size of 77.3 kbp (Fig. 2b, Suppl. Table 4). These regions overlapped with 11 gene models (Suppl. Table 5). Finally, the third analysis aimed at identifying associations between population allele frequencies and altitude through the computation of Bayes Factors (*BF*), and identified a single 7.43 kbp genomic interval located on chromosome 4 (Fig. 2c, Suppl. Table 6), which overlapped with two gene models (Suppl. Table 7). No significant gene ontology enrichment was detected in any of the three candidate gene sets. While there was only little overlap between *C*_2_ and *X^t^X* regions (13 genomic intervals), all three statistics detected the same major signal between 12 and 13 Mbp on chromosome 4 (Suppl. Fig. 2).

### A strong putative selective sweep on chromosome 4

To further dissect the signature of selection detected on chromosome 4, site-specific and multilocus windowed pairwise *F*_ST_ estimates were computed from pool-seq data within each locality to identify which population pairs were driving the signal. While little to no divergence was observed in the Spanish and Italian population pairs, divergence was accrued in the Mont Ventoux and Corsican pairs (Fig. 3). Interestingly, for these two pairs, this region was the most differentiated genome-wide between low and high altitude populations, albeit at different levels (*F*_ST_ estimates of 0.52 and 0.15 for Corsica and Mont Ventoux, respectively, Suppl. Fig. 3). Parallel divergence signatures between Corsican and Mont Ventoux population pairs could be caused by the introgression of an advantageous haplotype from one locality to the other. To test for this hypothesis, we computed the normalized *F*_4_ statistic (Patterson’s D, Green et al. 2010) along the genome using the poolfstat R package, testing all possible configurations between both Corsican and Mont Ventoux populations, Spanish populations and recently released North African *T. pityocampa* data as outgroups (Muller et al. 2025). Unfortunately, none of the tested configurations met the treeness criterion genome-wide, suggesting gene flow between populations and limiting our ability to detect putative adaptive introgression with the data at hand (data not shown). We thus focused on the Corsican pair to better characterize patterns of nucleotide diversity and divergence at this locus (Fig. 4, see Suppl. Fig. 4, 5 and 6 for results in the other localities).

**Figure 3.**
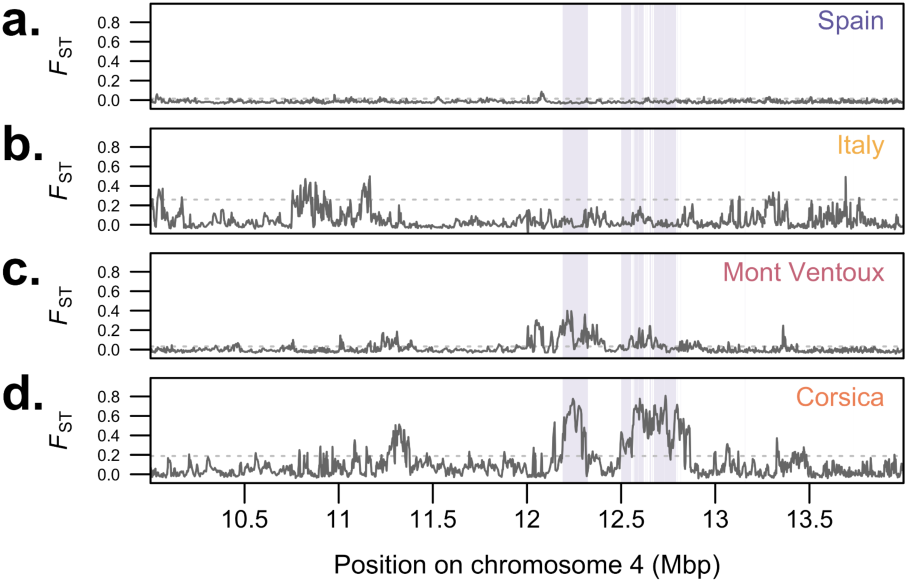
Pairwise *F*_ST_ divergence patterns within each locality in the main outlier genomic region on chromosome 4 (pool-seq data). Plain lines represent 100-SNP sliding window multilocus estimates. Purple blocks indicate candidate *X^t^X* regions identified with the local score approach. Dotted lines give the 95^th^ quantile of the genome-wide *F*_ST_ distribution for each population pair. **a.** Spain. **b.** Italy. **c.** Mont Ventoux. **d.** Corsica.

**Figure 4.**
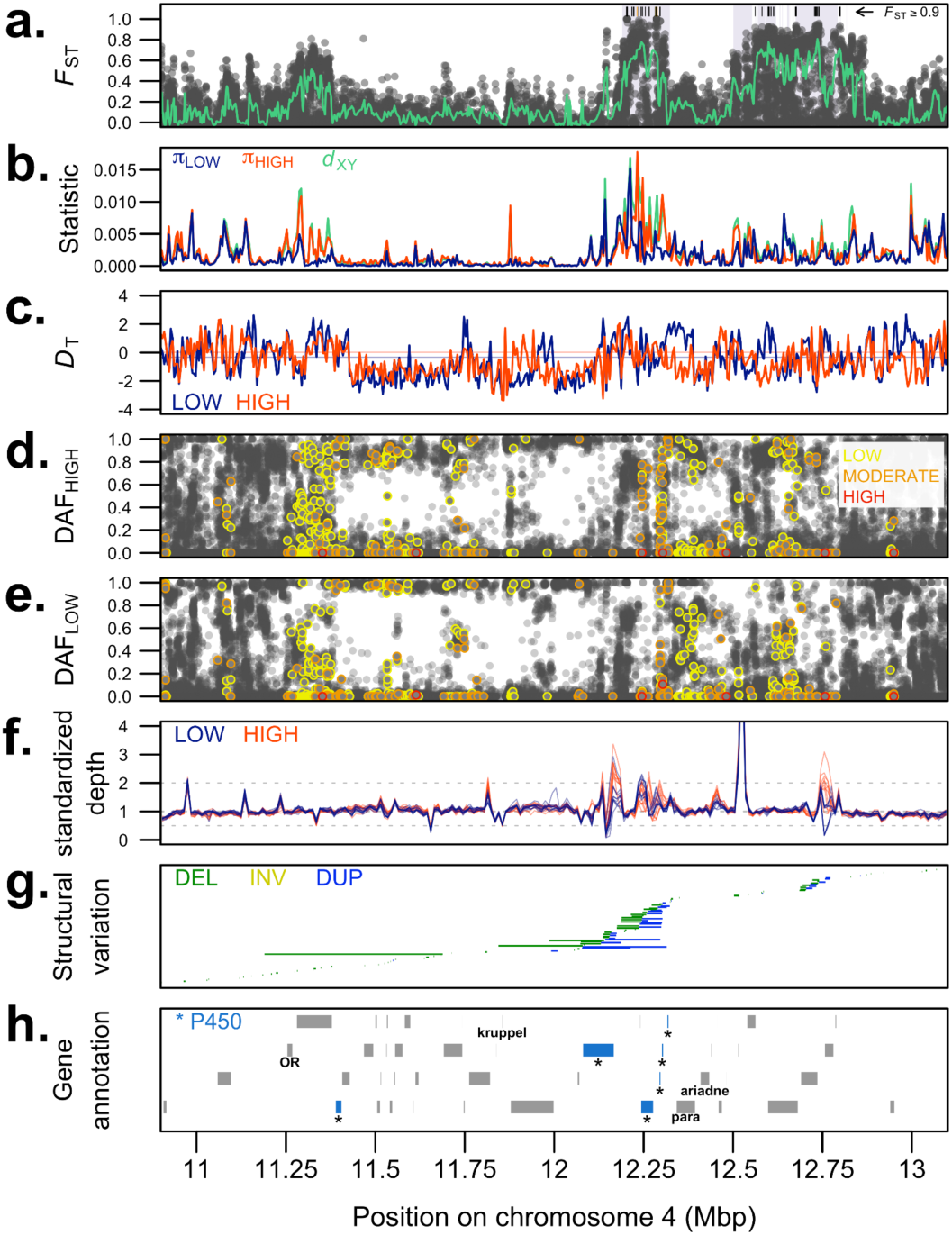
Zoom on the selective sweep signal detected between the two Corsican populations on chromosome 4. **a.** SNP-specific pairwise *F*_ST_ values between low and high altitude pools (green: multilocus *F*_ST_ values computed in sliding 100-SNP windows). Purple blocks indicate candidate *X^t^X* regions identified with the local score approach. **b.** Nucleotide diversity within (π) and absolute divergence between (*d*_XY_) populations, computed from ind-seq data in 5 kbp non-overlapping windows. **c.** Tajima’s *D* computed from pools in 5 kbp non-overlapping windows, with genome-wide means indicated as plain lines. **d.** Derived allele frequencies in high and **e.** low altitude pools, with SNPs colored according to their estimated functional impact predicted with SNPeff (yellow: synonymous, orange: non-synonymous, red: putative loss-of-function). **f.** Standardized sequencing depths computed from ind-seq libraries in 10 kbp windows. Dotted lines at 0.5 and 2 indicate halved and doubled average depths, respectively. **g.** Results of a screening for structural variation ≥ 500 basepairs using all Corsican ind-seq libraries jointly (DEL: deletion, INV: inversion, DUP: duplication). **h.** Gene annotation track (cytochrome P450 genes indicated in blue with asterisks).

Pairwise *F*_ST_ estimates between low and high Corsican pools were the highest between 12.2 and 12.8 Mbp, with 69 SNPs displaying *F*_ST_ values exceeding 0.9, including 14 intergenic SNPs alternatively fixed between low and high populations in the pool-seq data (Fig. 4a). Close to this region of elevated relative divergence, nucleotide diversity π and absolute divergence *d*_XY_ computed from ind-seq data were low in both populations between 11.375 and 12.13 Mbp, while flanking regions showed higher absolute divergence levels (Fig. 4b). In this same low nucleotide diversity region, Tajima’s *D* values were negative for both populations (Fig. 4c), which both showed deficits of SNPs at intermediate frequencies (Fig. 4d, Fig. 4e). Altogether, these observations (low nucleotide diversity and negative Tajima’s *D*) are in agreement with a scenario of selective sweep, and the functions of genes in the region may provide insights into the factors that have driven this putative selective event.

### Functional insights from the sweep region highlight detoxification mechanisms

Narrowing the focus to the 12-13 Mbp interval on chromosome 4, which encompasses the candidate *X^t^X* regions, we recovered 20 gene models, including a cluster of 5 cytochrome P450 monooxygenase genes from the *CYP6B* gene family (between 12.08 and 12.32 Mbp, with a sixth *CYP6B* gene outside the interval around 11.33 Mbp, see Fig. 4 and Suppl. Fig. 7 for visualisation, and Suppl. Table 8 for detailed statistics). Importantly, the expert annotation of detoxification genes in the pine processionary genome confirmed all P450 gene models predicted within this region were complete and showed alternatively spliced forms (Gautier et al. 2026), with expression support from RNA-seq data across multiple populations and developmental stages (Suppl. Fig. 8). Both individual sequencing depth profiles and short-read-based structural variation screening suggest a complex structural landscape in the vicinity of the cytochrome P450 cluster (Fig. 4f and 4g). Despite our stringent SNP filtering scheme, such indirect evidence for structural variation in the genomic region makes the fine mapping of putative causal substitutions within P450 genes hazardous. The 12-13 Mbp region also contains the voltage-gated sodium channel gene *para*, which is located 23 kbp away from the P450 cluster, in-between two candidate *X^t^X* regions (ca. 12.34 Mbp, Fig. 4 and Suppl. Fig. 7). Of note, no evidence of adaptive differentiation or altitude-associated variation was detected for SNPs within *para* (Suppl. Fig. 7, synonymous variants in yellow).

Among other genes in this 1 Mbp interval, an endonuclease-reverse transcriptase located only 3 kbp upstream of the P450 cluster showed strong signals of adaptive divergence (including one highly differentiated non-synonymous substitution, log_10_(*P_XtX_*) = 12.0, *F*_ST_ ^Corsica^ = 0.92). However, the high number of non-synonymous substitutions observed within this gene (Suppl. Fig. 7) suggests that it may be undergoing pseudogenization. To investigate this possibility, we assessed expression support using stage-specific transcriptomic datasets originally generated for the functional annotation of the genome (adults, pupae, larvae, and eggs from three distinct populations; see Gautier et al. 2026). The consistently low expression levels observed across multiple populations and developmental stages are consistent with pseudogenization, although tissue- or cell-type-specific expression patterns not captured in the current dataset cannot be excluded (Suppl. Fig. 8). Finally, a group of four genes located between 12.6 and 12.8 Mbp also displayed moderate *X^t^X* signatures and strong signals of differentiation between the Corsican population pairs. These four genes included two oxidoreductase enzymes (LOC113503460), along with homologs of *Rabconnectin-3A* and *Tempura*, both of which are involved in the Notch signaling pathway in *Drosophila* (Charng et al. 2014; Tuttle et al. 2014).

## Discussion

Our replicated genome-wide scan across four altitudinal gradients in the pine processionary moth uncovered a strong and repeated footprint of selection across a 1.5 Mb region located on chromosome 4. With a marked reduction in nucleotide diversity, negative Tajima’s *D*, and near-fixation of alternative alleles between Corsican populations, this region displays the classical genomic hallmarks of a recent selective sweep. Importantly, no functional assays have been conducted to confirm the causal role of the candidate genes identified here, and the interpretations proposed below regarding the putative selective agent(s) remain hypothesis-generating rather than confirmatory. While the same genomic region shows elevated, albeit less extreme, divergence levels between populations from Mont Ventoux (Suppl. Fig. 5), the magnitude and fine-scale pairwise *F*_ST_ profile of elevation-associated divergence differ among both localities, suggesting only partial genomic parallelism (Suppl. Fig. 7). Given the lack of evidence for recent introgression between these localities and the strong geographic structure in the dataset, this pattern may reflect partially independent local evolutionary trajectories acting on the same functional locus or pathway and/or spatial heterogeneity in the selective pressures experienced by populations (Battey et al. 2020). However, with the environmental data at hand, the selective agent underlying this pattern remains uncertain. In this context, we discuss below how differences in insecticide exposure, host-plant chemistry, or other ecological factors may all contribute to variation in the strength and genomic architecture of the response.

### A strong selective sweep signature suggestive of insecticide resistance?

Recurrent, independent sweeps at the same locus have been documented in several insect pests evolving resistance through gene amplification, regulatory changes, or target-site substitutions (Bass and Field 2011; Pélissié et al. 2022; Chen et al. 2025). Remarkably, the putative sweep encompasses a cluster of *CYP6B* cytochrome P450 monooxygenase genes and the voltage-gated sodium channel gene *para*, both widely implicated in insecticide detoxification and target-site resistance in insects (Nelson et al. 1996; Puinean et al. 2010; Liu 2015; Kaduskar et al. 2022; Pu and Chung 2024; Ngegba et al. 2025). While there is little support in our dataset for point mutations driving reduced sensitivity to insecticides in the *para* gene, coverage profiles and short-read structural-variant scans reveal a complex structural landscape surrounding the *CYP6B* cluster, potentially including duplications and/or rearrangements. Copy number expansion of P450 genes is a well-characterised mechanism leading to xenobiotic resistance, where additional copies of detoxification genes increase transcriptional output or provide substrates for neofunctionalization (Scott 1999; Bass and Field 2011). Moreover, recent mechanistic studies in *Spodoptera exigua* showed that both cis- and trans-regulatory changes synergistically increase P450 expression to confer resistance to both organophosphate and pyrethroid insecticides (B. Hu et al. 2025), suggesting that both structural and regulatory evolution could drive the putative selective sweep detected in the vicinity of the *CYP6B* cluster.

The role of *CYP6* genes in pesticide detoxification and resistance is well established (Nauen et al. 2022). In Lepidoptera, their overexpression is classically associated with resistance to synthetic pyrethroids (e.g., Bautista et al. 2009; Xu et al. 2020; Perini et al. 2021; do Nascimento et al. 2023). Pyrethroids act as neurotoxic agents that disrupt the insect’s nervous system, mostly targeting voltage-gated sodium channels in insect nerve cells, causing paralysis and death. In the European Union, forestry management remains largely under Member State jurisdiction, which results in spatial heterogeneity in treatment intensity and pest management practices (Leroy 2024). This heterogeneity, together with possible local derogations for banned compounds and undocumented applications, makes it difficult to reconstruct precise exposure histories for each sampling locality. Over the last three decades, pyrethroid use has progressively declined and has increasingly been replaced by *Bacillus thuringiensis* extracts (*Bt* toxin) at large scales or on caterpillars traps at local scales (Gui-ming et al. 2001). As *CYP6* genes have so far not been shown to be implicated in resistance to *Bt* toxin (Heckel et al. 2007; D. Hu et al. 2025), and under the insecticide exposure hypothesis, the putative selective sweep identified likely predates the adoption of *Bt*-based control and could have been caused by the previous use of pyrethroids. The maintenance of resistance alleles in natural populations following the ban on pyrethroids could be explained primarily by the relatively recent effective ban of these compounds and/or by a potentially low cost of the resistance mechanism.

Interestingly, Calla et al. (2021) investigated insecticide resistance in another moth, the navel orangeworm *Amyelois transitella*, which is a major almond pest in US orchards. Comparing populations with different levels of resistance to bifenthrin, a pyrethroid, they also detected a putative hard selective sweep covering 1.3 Mbp and encompassing the same candidate P450 cluster and *para* gene, suggesting gene order conservation between *A. transitella* and *T. pityocampa* despite >100 MY of divergence (the syntenic block extends beyond *kruppel* and *ariadne*, shown on Fig. 4h). In the future, long-read data combined with pangenomic approaches could help characterize the structural complexity of this genomic region (Secomandi et al. 2025), confirm the presence of structural variants, and determine their distribution among pine processionary populations with different exposure histories. Such approaches would also help resolve copy number variation (Puinean et al. 2010), transposable element content (Baril et al. 2023) and haplotype structure at this locus, which cannot be reliably inferred from short-read data alone. In parallel, gene expression profiling across populations and/or controlled exposure experiments under different insecticide treatments could help assess the phenotypic and regulatory consequences associated with the variation we identified (see, e.g., Wondji et al. 2022).

### Tree secondary metabolites as potential drivers of detoxification genes?

Another potential selective pressure driving P450 expansion is resistance to natural tree defense compounds such as α-pinene, a monoterpene present in pine tissues, including the needles that caterpillars feed on (Kubeczka and Schultze 1987). P450s (notably *CYP4* and *CYP6* gene families) are central to pinene detoxification in Coleoptera such as bark beetles (Dai et al. 2014; Dai et al. 2015; Chiu et al. 2019) and longhorn beetles (He et al. 2022). In these species, the expression of detoxification genes in response to α-pinene and other monoterpenes has been shown to vary among populations exposed to different host trees or terpene concentrations (He et al. 2022; Torres-Banda et al. 2022). While evidence is less abundant in Lepidoptera, α-pinene has been shown to induce overexpression of *CYP6B2* in the cotton bollworm *Helicoverpa armigera* (Ranasinghe et al. 1997), and exposure to α-pinene led to both oxidative stress and nervous system disruption in the flour moth *Ephestia kuehniella* (Shahriari et al. 2018), increasing the enzymatic activity of superoxide dismutases and glutathione S-transferases (P450s were not assayed in the study). Interestingly, the Corsican populations between which we detected the strongest putative sweep signal were sampled on different host trees, *P. pinaster* at low elevation and *P. nigra* var. *corsicana* (syn*. P. nigra* ssp. *laricio*) at high elevation (Table 1). These pine species differ markedly in their needle essential oil composition, including α-pinene content (Ioannou et al. 2014). raising the possibility that variation in host-associated chemical environments could contribute to the selective pattern we detected. Under this hypothesis, differences in terpene exposure among host trees may exert selection on detoxification pathways involving the CYP6 cluster identified here. However, because present-day host trees may not accurately reflect the historical host environment experienced by these populations, terpene composition and concentration can vary substantially between individuals within species depending on both biotic and abiotic factors (Kopaczyk et al. 2020), and because multiple selective pressures may act simultaneously on detoxification genes, this interpretation remains speculative. Further ecological and experimental investigations (including, e.g., transcriptomic assays under controlled conditions) will therefore be required to disentangle the respective contributions of host-plant chemistry and anthropogenic xenobiotic exposure to the evolution of detoxification mechanisms in the pine processionary moth.

### Weak signatures of altitudinal adaptation in the pine processionary moth

The genomics of altitudinal adaptation are increasingly studied in Lepidoptera, revealing both parallel and polygenic genomic architectures. At a larger evolutionary timescale compared to our study, genome-wide scans in *Heliconius erato* and *H. melpomene* along replicated altitudinal transects identified multiple divergent regions associated with altitude, showing that adaptation often draws on standing variation or adaptive introgression from related taxa and involves moderate-effect loci distributed across the genome (Montejo-Kovacevich et al. 2022). In the same system, a related genome-wide association study of wing aspect ratio showed that altitude-related morphological shifts are heritable and highly polygenic (Montejo-Kovacevich et al. 2019). In our study, outside the major signal on chromosome 4, we found only a few genomic regions displaying elevated levels of divergence between low- and high-altitude groups of populations and/or association with altitude (*C*_2_ and *BF* statistics, respectively), for which gene content analysis could not provide any clear link between gene function and selective pressures possibly experienced in the wild (see Suppl. Tables 3, 5 and 7). Of note, while the *X^t^X* analysis aimed at detecting local adaptation signatures uncovered 721 genomic regions totaling 1.82 Mbp, the *C*_2_ analysis only identified 23 genomic regions totaling 77.3 kbp, thus suggesting that altitude may not exert a major selection pressure, at least within our study area. Alternatively, assuming altitude is a major stressor for the pine processionary moth, such sparse signals can emerge under several non-mutually exclusive scenarios, including polygenic adaptation governed by many small-effect loci (Pritchard et al. 2010), substantial phenotypic plasticity buffering selection (Forsman 2015), heterogeneous responses to selection across localities (Barghi et al. 2020), or interference from stronger, unrelated selective forces (McCulloch and Waters 2023). Adaptation to altitude is also influenced by a combination of factors, including pine processionary phenology (Martin et al. 2022), temperature, air pressure differences, and changes in habitat structure and composition (Sømme 1989). Adding to this complexity, organismal responses to these factors can be correlated, and disentangling their individual effects from our observational data alone is challenging. While the methods we used to recover divergence and association signals from genome-wide polymorphism data show modest to good performance to detect polygenic adaptation in simulations (Gautier 2015), pinpointing small-effect loci remains a difficult task (Barghi et al. 2020). Additionally, altitudes were not fully balanced between localities in our sampling scheme (Table 1), owing to difficulties in localizing and/or sampling populations and/or nests, and such imbalance could hamper the power of our association analyses (e.g., the elevation of the low-elevation Spanish population is higher than that of the high-elevation Corsican population).

### Tangled selective pressures in natural populations

Our study exemplifies the difficulty of teasing apart ecological and anthropogenic selection in natural populations. While our results suggest altitude may not exert a major selection pressure in our studied populations, in many arthropods, xenobiotic resistance evolves orders of magnitude faster than climatic adaptation because pesticide exposure imposes extreme and episodic selective pressures (McCulloch and Waters 2023). Such responses may evolve more rapidly in phytophagous insects, in which insecticide resistance mechanisms can rely partly on genes, such as cytochrome P450s, which are already involved in the metabolism of plant defence compounds (preadaptation hypothesis, Gordon 1961; Croft et al. 1983; Rosenheim et al. 1996; Dermauw et al. 2013). In addition, linkage and pleiotropy might blur these processes: selection on detoxification genes could indirectly alter physiological traits related to thermal or oxidative stress, while climate-driven selection could modify resistance phenotypes (e.g., Rostant et al. 2017). Spatial heterogeneity exposure further complicates inference (Battey et al. 2020). Localized pesticide use can generate parallel sweeps in distant populations facing similar treatment regimes but leave other populations unaffected, creating an apparent mismatch between ecological and genomic signals. Disentangling these effects requires a wealth of environmental data, along with controlled assays, which are not realistic in most settings. Repeated sampling of the same localities in the future could nevertheless provide valuable insight into the temporal stability of the observed genomic patterns under ongoing environmental change. However, this work also shows that careful interpretation of genomic data can point towards unexpected selection pressures. From a management perspective, and under the insecticide exposure hypothesis, the potential for resistance evolution in *T. pityocampa*, which is in theory not subject to routine treatment, highlights the need for caution and control in chemical interventions carried in natural environments.

## Methods

### Study populations and samples

Sampling was performed between October 2011 and March 2014 for four pairs of populations along altitudinal gradients located in the Spanish Sierra Nevada, the Italian Alps, Corsica and Mont Ventoux (France; Table 1 and Figure 1). For each of the four pairs, individuals were sampled for both an upper slope population and a lower slope population (altitudes are given in Table 1). For Corsican and Italian populations, larvae were directly collected from the nests during winters between 2011 and 2014 (Table 1). For Spain and Mont Ventoux, male adults were sampled using pheromone traps during summers 2012 and 2013 (Table 1). These traps were baited with (Z)-13-hexadecen-11-ynyl acetate (“pityolure”), a specific attractant which is the only compound present in female pheromone glands (Guerrero et al. 1981; Quero et al. 1997). In the core of the species’ distribution, including the localities sampled in the present study, pheromone trapping has been shown to accurately capture the genetic diversity present within populations (Salvato et al. 2005). All individuals were preserved in 95% ethanol after sampling. To decrease relatedness among individuals both within populations and within pools (see below), a single caterpillar per colony and per tree was collected for both Italian and Corsican populations, while for pheromone traps, individuals were collected using several traps deployed over most of the emergence period (six to eight weeks).

### Lab work and whole-genome sequencing

DNA from each individual was extracted using Qiagen’s commercial DNeasy Blood & Tissue Kit. Population samples ranged from 26 to 40 individuals per population (Table 1). For pooled sequencing, DNA extractions were mixed in equimolar proportions for each population. For individual sequencing, the DNA of 8 to 16 individuals per population were selected, for a total of 89 individuals. A total of 97 libraries (89 ind-seq + eight pool-seq) were constructed using Illumina’s TruSeq nano DNA kit following manufacturer instructions. Paired-end sequencing was carried out at the Montpellier GenomiX (MGX) platform on Illumina HiSeq 2500 (pool-seq samples, read length: 125 nucleotides) and Illumina NovaSeq 6000 (ind-seq samples, read length: 150 nucleotides), targeting mean read depths of 60× and 20× for pool-seq and ind-seq libraries, respectively. Sequencing produced between 92.3-682 (median: 123) and 260-577 (median: 374) million reads for ind-seq and pool-seq libraries, respectively, with mean insert sizes ranging from 340 to 457 (median: 387). Mean depths ranged from 15.7× to 109× (median: 21.9×) for ind-seq libraries and from 41.4× to 97.7× (median: 60.4×) for pool-seq libraries (see Suppl. Table 1 for detailed sequencing statistics).

### Bioinformatics

Unless stated otherwise, all analyses and data visualization were carried in R v4.5.1 (R Core Team 2019). Data quality was checked with FastQC v0.12.0 (Andrews 2010) before and after trimming of raw reads using fastp v0.23.2 (Chen et al. 2018). Trimmed reads were aligned against the chromosome-level pine processionary genome assembly Tpit_2.1 (Gautier et al. 2026) using bwa mem v0.7.18 (Li 2013) and duplicates were removed with sambamba v1.0.1 (Tarasov et al. 2015). Insert sizes were collected from BAM files using Picard v3.0.0 (http://broadinstitute.github.io/picard), and sequencing depths were tallied for each library in 10 kbp windows along the genome with mosdepth v0.3.6 (Pedersen and Quinlan 2018). SNP calling was performed jointly on all ind-seq and pool-seq libraries using the haplotype-aware caller FreeBayes v1.3.6 (Garrison and Marth 2012) with the following parameters: `--pooled-continuous -m 20 -q 20 -p 2 -n 3 --min-coverage 10 --limit-coverage 300 -E -1 --max-complex-gap - 1 --haplotype-length -1`. Multi-nucleotide variants were decomposed with vt v0.57721 (Tan et al. 2015), and only biallelic sites displaying qualities ≥ 30 and supported by at least one forward and one reverse reads across all ind-seq and pool-seq libraries were retained using BCFtools v1.17 (Danecek et al. 2021).

The resulting VCF file was split into pool-seq (*n* = 8 samples) and ind-seq VCF files (*n* = 89 samples) to apply library-specific filtering schemes. For the pool-seq VCF file, sites were filtered using the poolfstat v3.0.0 R package (Gautier et al. 2022) to retain SNPs displaying overall minor allele frequencies (MAF) computed on read counts ≥ 0.05, at least four observations of the minor allele across all pools, sequencing depths comprised between 10× and 200× in each pool, and discarding sites located at less than five basepairs from an indel. For the ind-seq VCF file, since samples contained larvae for which sex cannot be determined based on morphology, Z-chromosome-to-autosomal sequencing depth ratios were computed for each ind-seq sample. Females are heterogametic in Lepidoptera and should display sequencing depth ratios around 0.5 (one Z copy for two autosomal copies), compared to unity for males. The depth ratio analysis identified 66 males and 23 females from the ind-seq data, with all individuals sampled using pheromone traps assigned as males, as expected (Table 1, Suppl. Table 1, Suppl. Fig. 1). After this bioinformatic sex determination step, biallelic sites displaying depths below 8× and above twice the mean library depth for each individual were coded as missing using BCFtools. The lower filter was adjusted for Z-linked sites in females (allowing minimum 4×). Sites displaying more than 25% missing data across all individuals were discarded, along with sites which did not fulfil the global MAF ≥ 0.05 criterion (computed across all ind-seq libraries). Overall, joint SNP calling and subsequent library-specific filtering recovered 14,323,239 sites for the pool-seq dataset and a subset of 9,193,257 sites for the ind-seq dataset. Additionally, in order to compute unbiased nucleotide diversity and divergence metrics, a VCF file containing both variant and invariant sites (hereafter, all sites) was generated solely from ind-seq data using BCFtools, following recommendations from Korunes & Samuk (2021). Below, distinct analyses were conducted on the ind-seq and pool-seq datasets using methods specifically adapted to each data type.

### Population structure, divergence and nucleotide diversity metrics

Population structure was first documented from ind-seq data using a principal component analysis (PCA) run on 84,443 linkage-pruned autosomal genotypes with plink v2.0 (pruning carried with parameters --indep-pairwise 250kb 1 0.2, Chang et al. 2015). Additionally, unsupervised clustering was carried on the same dataset with Admixture v1.3.0 (Alexander et al. 2009) using default parameters and ranging ancestry components *K* from 2 to 12, with 10 replicates per *K* value. Results were aggregated with StructureSelector (Li and Liu 2018) and the most likely number of ancestry components was estimated following Puechmaille (2016), owing to unbalanced sample sizes across populations. Nucleotide diversity (π) and pairwise divergence metrics (*F*_ST_ and *d*_XY_) were computed from the all sites ind-seq VCF file in 5 kbp windows along the genome using pixy v1.2.11 (Korunes and Samuk 2021).

### Inference of demographic history through time

We used the sequential Markovian coalescent model implemented in MSMC v2.1.4 (Schiffels and Wang 2020) to infer historical changes in effective population sizes (*N*_e_), using the three most covered individuals per population (SNP depth > 23×). Phasing and imputation were carried jointly for all 24 ind-seq samples (3 individuals × 8 populations) with Beagle v5.4 (Browning et al. 2021), using default parameters. For each individual, callable sites were tallied using mosdepth v0.3.6 (Pedersen and Quinlan 2018) using the same depth parameters as for the SNP calling step (see above). These intervals were combined with the repeat GFF file from Gautier et al. (2026) to mask sites. MSMC2 was then run within each population (6 haplotypes) following Schiffels and Wang (2020) using 19 epochs to limit overfitting (`-p 1*2+15*1+1*2`). Results were visualised assuming a mutation rate *µ* = 2.9 × 10^-9^ (Keightley et al. 2015) and a single generation per year.

### Identification of candidate genomic regions associated with altitude

Genomic regions displaying altitudinal variations in allele frequencies were identified using the hierarchical Bayesian framework implemented in BayPass v3.0.0 (Gautier 2015), which explicitly accounts for the covariance structure among population allele frequencies resulting from the shared history of samples. Three distinct analyses were carried out for each SNP of the pool-seq dataset under BayPass’ standard covariate model, specifying pool haploid sample sizes (assuming all individuals within a pool are male and carry two copies of the Z chromosome, which is conservative) and using default options for the importance sampling approximation. First, the *X^t^X* statistic (Günther and Coop 2013) was computed to detect signals of positive selection in any population, irrespective of elevation levels. Second, the *C*_2_ statistic (Olazcuaga et al. 2020) was computed to contrast allele frequencies between the low and high altitude groups of populations (i.e., altitude is treated as a binary trait, with two groups of four populations each). Third, Bayes factors (*BF*) were computed to measure the strength of evidence for the linear association between population allele frequencies and altitude in meters (scaled by default in BayPass, so that *μ* = 0 and *σ*^2^ = 1). For all three analyses, autosomal and Z-linked SNPs were treated separately, and to alleviate the computational burden, the full dataset was thinned into subsets containing ca. 75,000 SNPs each (leading to 185 and 6 subsets for autosomal and Z compartments, respectively). The local score approach developed by Fariello et al. (2017) was then applied to the statistics computed with BayPass to account for linkage. This approach implemented in BayPass’ companion R script combines statistics at neighboring SNPs with global MAF ≥ 10% (overall reference allele frequency estimated by BayPass) along each chromosome to detect significant regions based on the user-defined significance threshold ξ. To better prioritize candidate genomic regions, distinct thresholds were applied to the different statistics: ξ = 2 for both *C*_2_ and *BF* (significance level *α* = 0.01), versus ξ = 3 for *X^t^X* (*α* = 0.001). For each analysis, gene models located within candidate genomic regions were extracted from the genome annotation file and gene ontology enrichments were tested using the topGO R package v2.60.1 (Alexa and Rahnenfuhrer 2025).

### Fine-scale characterization of the main candidate selective sweep region

Genome-wide analyses using BayPass recovered a strong signal on chromosome 4 (see Results), which was further characterized at a finer scale. To identify which population pair(s) drove the signal in the genomic region, pairwise *F*_ST_ metrics were computed between low and high populations within each locality from pool-seq data. Then, focusing on the Corsican population pair, which showed the highest relative divergence in the region (see Results), and in addition to π and *d*_XY_ metrics previously computed from ind-seq data, Tajima’s *D* (*D*_T_) was computed from pool-seq data in 5 kbp windows using npstat v1 with default options (Ferretti et al. 2013). All SNPs passing previous filters in the pool-seq VCF file were polarised by setting as ancestral the major allele from the two Spanish populations, as Spain is assumed to be a major glacial refugium of Western European pine processionary populations (Kerdelhué et al. 2009). These polymorphisms were further annotated with SNPeff v4.0 (Cingolani 2022) using the Tpit_2.1 GFF annotation file (Gautier et al. 2026). Finally, to better characterize the structural landscape in the candidate genomic region, sequencing depths were tallied for each Corsican ind-seq library in 10 kbp windows using mosdepth v0.3.6 (Pedersen and Quinlan 2018). Additionally, evidence of structural variation (SV) was gathered from the same dataset in the region using smoove v0.2.8 (https://github.com/brentp/smoove), a wrapper which implements internal read filtering steps and combines the Lumpy SV caller (Layer et al. 2014) to Duphold for depth-based SV annotation (Pedersen and Quinlan 2019). Detecting structural variation from short-read data is a complex endeavor (see, e.g., Olson et al. 2023), and results from this last analysis should thus be interpreted cautiously.

## Data accessibility

The data underlying this article have been deposited in the European Nucleotide Archive (ENA) at EMBL-EBI under accession number PRJEB114432 (https://www.ebi.ac.uk/ena/browser/view/PRJEB114432). Sampling locations are provided in Table 1 and sample metadata are provided in Suppl. Table 1.

## Acknowledgments

This work was funded through both the ANR ANR-22-CE02-0009 LOADEXP and INRAE exploratory projects to C.P. and ANR-10-JCJC-1705-01 GENOPHENO to C.K. We are grateful to Aurélie Ducasse, Renaud Vitalis, Vincent Danneville, Jean-Michel Gigleux, Pikria Doinjashvili and Cécile Robin for their assistance with project management. We acknowledge our colleagues Andrea Battisti (Univ. Padova, Italy), José A. Hódar and Rut Aspizua (Univ. Granada, Spain), Jérôme Rousselet, Alain Roques and Francis Goussard (URZF, INRAE Orléans Val de Loire, France) and Jean-Claude Martin, Anne-Sophie Brinquin, Marianne Correard, René Mazet and Estelle Morel (UEFM, INRAE PACA, France) for providing samples from Italy, Spain, Corsica and Mont Ventoux, respectively. Sequencing services were provided by the MGX facility (https://mgx.cnrs.fr). We acknowledge the GenSeq platform (Université Montpellier) for granting us access to their facilities for library preparation. We are grateful to the genotoul bioinformatics platform Toulouse Occitanie (Bioinfo Genotoul, https://doi.org/10.15454/1.5572369328961167E12) and to the CBGP cluster (Nathalie Vieira and Sylvain Piry) for providing computing and storage resources.

## Bibliography

Alexa A, Rahnenfuhrer J. 2025. topGO: Enrichment Analysis for Gene Ontology. Available from: https://bioconductor.org/packages/topGO

Alexander DH, Novembre J, Lange K. 2009. Fast model-based estimation of ancestry in unrelated individuals. Genome Res. 19:1655–1664.

Andrews S. 2010. FastQC: a quality control tool for high throughput sequence data. Available from: https://www.bioinformatics.babraham.ac.uk/projects/fastqc

Barghi N, Hermisson J, Schlötterer C. 2020. Polygenic adaptation: a unifying framework to understand positive selection. Nat. Rev. Genet. 21:769–781.

Baril T, Pym A, Bass C, Hayward A. 2023. Transposon accumulation at xenobiotic gene family loci in aphids. Genome Res. 33:1718–1733.

Bass C, Field LM. 2011. Gene amplification and insecticide resistance. Pest Manag. Sci. 67:886–890.

Battey CJ, Ralph PL, Kern AD. 2020. Space is the place: Effects of continuous spatial structure on analysis of population genetic data. Genetics 215:193–214.

Battisti A, et al. 2005. Expansion of geographic range in the pine processionary moth caused by increased winter temperatures. Ecol. Appl. 15:2084–2096.

Bautista MAM, Miyata T, Miura K, Tanaka T. 2009. RNA interference-mediated knockdown of a cytochrome P450, CYP6BG1, from the diamondback moth, Plutella xylostella, reduces larval resistance to permethrin. Insect Biochem. Mol. Biol. 39:38–46.

Bellard C, Bertelsmeier C, Leadley P, Thuiller W, Courchamp F. 2012. Impacts of climate change on the future of biodiversity: Biodiversity and climate change. Ecol. Lett. 15:365–377.

Bernatchez L, Ferchaud A-L, Berger CS, Venney CJ, Xuereb A. 2024. Genomics for monitoring and understanding species responses to global climate change. Nat. Rev. Genet. 25:165–183.

Bonhomme M, et al. 2010. Detecting selection in population trees: The Lewontin and Krakauer test extended. Genetics 186:241–262.

Bourgeois YXC, Warren BH. 2021. An overview of current population genomics methods for the analysis of whole-genome resequencing data in eukaryotes. Mol. Ecol. 30:6036– 6071.

Bourougaaoui A, Ben Jamâa ML, Robinet C. 2021. Has North Africa turned too warm for a Mediterranean forest pest because of climate change? Clim. Change 165:1–20.

Browning BL, Tian X, Zhou Y, Browning SR. 2021. Fast two-stage phasing of large-scale sequence data. Am. J. Hum. Genet. 108:1880–1890.

Calla B, et al. 2021. Selective sweeps in a nutshell: The genomic footprint of rapid insecticide resistance evolution in the almond agroecosystem. Genome Biol. Evol. 13:evaa234.

Chang CC, et al. 2015. Second-generation PLINK: rising to the challenge of larger and richer datasets. Gigascience 4:7.

Charng W-L, et al. 2014. Drosophila Tempura, a novel protein prenyltransferase α subunit, regulates notch signaling via Rab1 and Rab11. PLoS Biol. 12:e1001777.

Chen I-C, Hill JK, Ohlemüller R, Roy DB, Thomas CD. 2011. Rapid range shifts of species associated with high levels of climate warming. Science 333:1024–1026.

Chen L, et al. 2025. Population genomics reveal multiple independent origins of pesticide resistance in the polyphagous pest, Tetranychus urticae. Commun. Biol. 8:1197.

Chen S, Zhou Y, Chen Y, Gu J. 2018. fastp: an ultra-fast all-in-one FASTQ preprocessor. Bioinformatics 34:i884–i890.

Chiu CC, Keeling CI, Bohlmann J. 2019. The cytochrome P450 CYP6DE1 catalyzes the conversion of α-pinene into the mountain pine beetle aggregation pheromone trans-verbenol. Sci. Rep. 9:1477.

Cingolani P. 2022. Variant annotation and functional prediction: SnpEff. Methods Mol. Biol. 2493:289–314.

Croft BA, Strickler K, Georghiou GP, Saito T. 1983. Pest resistance to pesticides.

Dai L, et al. 2015. Cytochrome P450s from the Chinese white pine beetle, Dendroctonus armandi (Curculionidae: Scolytinae): Expression profiles of different stages and responses to host allelochemicals. Insect Biochem. Mol. Biol. 65:35–46.

Dai L, et al. 2014. Two CYP4 genes of the Chinese white pine beetle, Dendroctonus armandi (Curculionidae: Scolytinae), and their transcript levels under different development stages and treatments: Dendroctonus armandi. Insect Mol. Biol. 23:598– 610.

Danecek P, et al. 2021. Twelve years of SAMtools and BCFtools. Gigascience [Internet] 10. Available from: 10.1093/gigascience/giab008

Dermauw W, et al. 2013. A link between host plant adaptation and pesticide resistance in the polyphagous spider mite Tetranychus urticae. Proc. Natl. Acad. Sci. U. S. A. 110:E113–E122.

Dudaniec RY, Yong CJ, Lancaster LT, Svensson EI, Hansson B. 2018. Signatures of local adaptation along environmental gradients in a range-expanding damselfly (Ischnura elegans). Mol. Ecol. 27:2576–2593.

Fariello MI, et al. 2017. Accounting for linkage disequilibrium in genome scans for selection without individual genotypes: The local score approach. Mol. Ecol. 26:3700–3714.

Ferretti L, Ramos-Onsins SE, Pérez-Enciso M. 2013. Population genomics from pool sequencing. Mol. Ecol. 22:5561–5576.

Forester BR, Lasky JR, Wagner HH, Urban DL. 2018. Comparing methods for detecting multilocus adaptation with multivariate genotype-environment associations. Mol. Ecol. 27:2215–2233.

Forsman A. 2015. Rethinking phenotypic plasticity and its consequences for individuals, populations and species. Heredity 115:276–284.

Futschik A, Schlötterer C. 2010. The next generation of molecular markers from massively parallel sequencing of pooled DNA samples. Genetics 186:207–218.

Garrison E, Marth G. 2012. Haplotype-based variant detection from short-read sequencing. arXiv [q-bio.GN] [Internet]. Available from: http://arxiv.org/abs/1207.3907

Gautier M. 2015. Genome-Wide Scan for Adaptive Divergence and Association with Population-Specific Covariates. Genetics 201:1555–1579.

Gautier M, et al. 2026. A Chromosome-Level Assembly of the Pine Processionary Moth (*Thaumetopoea pityocampa*) genome. J. Hered.:esag035.

Gautier M, Vitalis R, Flori L, Estoup A. 2022. ƒ-statistics estimation and admixture graph construction with Pool-Seq or allele count data using the R package poolfstat. Mol. Ecol. Resour. 22:1394–1416.

Gordon HT. 1961. Nutritional factors in insect resistance to chemicals. Annu. Rev. Entomol. 6:27–54.

Green RE, et al. 2010. A draft sequence of the Neandertal genome. Science 328:710–722.

Guerrero A, et al. 1981. Identification of a potential sex pheromone of the processionary moth, Thaumetopea pityocampa (lepidoptera, notodontidae). Tetrahedron Lett. 22:2013–2016.

Gui-ming L, Xiang-yue Z, Lu-quan W. 2001. The use ofBacillus thuringiensis on Forest Integrated Pest Management. J. For. Res. 12:51–54.

Günther T, Coop G. 2013. Robust identification of local adaptation from allele frequencies. Genetics 195:205–220.

Heckel DG, et al. 2007. The diversity of Bt resistance genes in species of Lepidoptera. J. Invertebr. Pathol. 95:192–197.

He X, et al. 2022. Transcriptome profiling and RNA interference reveals relevant detoxification genes in *Monochamus alternatus* response to (+)-α-pinene. J. Appl. Entomol. 146:823–837.

Hill JK, Griffiths HM, Thomas CD. 2011. Climate change and evolutionary adaptations at species’ range margins. Annu. Rev. Entomol. 56:143–159.

Hu B, et al. 2025. Two independent regulatory mechanisms synergistically contribute to P450-mediated insecticide resistance in a lepidopteran pest, Spodoptera exigua. BMC Biol. 23:122.

Hu D, Wang D, Pan H, Liu X. 2025. Molecular mechanisms underlying resistance to Bacillus thuringiensis Cry toxins in Lepidopteran pests: An updated research perspective. Agronomy 15:155.

Inouye DW. 2022. Climate change and phenology. Wiley Interdiscip. Rev. Clim. Change [Internet] 13. Available from: 10.1002/wcc.764

Ioannou E, Koutsaviti A, Tzakou O, Roussis V. 2014. The genus Pinus: a comparative study on the needle essential oil composition of 46 pine species. Phytochem. Rev. 13:741– 768.

Kaduskar B, et al. 2022. Reversing insecticide resistance with allelic-drive in Drosophila melanogaster. Nat. Commun. 13:291.

Kanat M, Alma MH, Sivrikaya F. 2005. Effect of defoliation by Thaumetopoea pityocampa (Den. & Schiff.) (Lepidoptera: Thaumetopoeidae) on annual diameter increment of Pinus brutia Ten. in Turkey. Ann. For. Sci. 62:91–94.

Keightley PD, et al. 2015. Estimation of the spontaneous mutation rate in Heliconius melpomene. Mol. Biol. Evol. 32:239–243.

Kerdelhué C, et al. 2009. Quaternary history and contemporary patterns in a currently expanding species. BMC Evol. Biol. 9:220.

Kopaczyk JM, Warguła J, Jelonek T. 2020. The variability of terpenes in conifers under developmental and environmental stimuli. Environ. Exp. Bot. 180:104197.

Korunes KL, Samuk K. 2021. pixy: Unbiased estimation of nucleotide diversity and divergence in the presence of missing data. Mol. Ecol. Resour. 21:1359–1368.

Kubeczka K-H, Schultze W. 1987. Biology and chemistry of conifer oils. Flavour Fragr. J. 2:137–148.

Layer RM, Chiang C, Quinlan AR, Hall IM. 2014. LUMPY: a probabilistic framework for structural variant discovery. Genome Biol. 15:R84.

Leroy BML. 2024. Global insights on insecticide use in forest systems: Current use, impacts and perspectives in a changing world. Curr. For. Rep. 11:6.

Li H. 2013. Aligning sequence reads, clone sequences and assembly contigs with BWA-MEM. arXiv [q-bio.GN] [Internet]. Available from: http://arxiv.org/abs/1303.3997

Liu N. 2015. Insecticide resistance in mosquitoes: impact, mechanisms, and research directions. Annu. Rev. Entomol. 60:537–559.

Li Y-L, Liu J-X. 2018. StructureSelector: A web-based software to select and visualize the optimal number of clusters using multiple methods. Mol. Ecol. Resour. 18:176–177.

Lotterhos KE, Whitlock MC. 2015. The relative power of genome scans to detect local adaptation depends on sampling design and statistical method. Mol. Ecol. 24:1031– 1046.

Martin J-C, Mesmin X, Buradino M, Rossi J-P, Kerdelhué C. 2022. Complex drivers of phenology in the pine processionary moth: Lessons from the past. Agric. For. Entomol. 24:247–259.

Martin J-C, Rossi J-P, Buradino M, Kerdelhué C. 2021. Monitoring of adult emergence in the pine processionary moth between 1970 and 1984 in Mont Ventoux, France. Biodivers. Data J. 9:e61086.

McCulloch GA, Waters JM. 2023. Rapid adaptation in a fast-changing world: Emerging insights from insect genomics. Glob. Chang. Biol. 29:943–954.

Montejo-Kovacevich G, et al. 2022. Repeated genetic adaptation to altitude in two tropical butterflies. Nat. Commun. 13:4676.

Montejo-Kovacevich G, et al. 2019. Altitude and life-history shape the evolution of Heliconius wings. Evolution 73:2436–2450.

Muller T, et al. 2025. Population genomics of incipient allochronic divergence in the Pine Processionary Moth. Mol. Ecol., 34: e70189.

do Nascimento ARB, et al. 2023. Susceptibility monitoring and comparative gene expression of susceptible and resistant strains of Spodoptera frugiperda to lambda-cyhalothrin and chlorpyrifos. Pest Manag. Sci. 79:2206–2219.

Nauen R, Bass C, Feyereisen R, Vontas J. 2022. The role of cytochrome P450s in insect toxicology and resistance. Annu. Rev. Entomol. 67:105–124.

Nelson DR, et al. 1996. P450 superfamily: update on new sequences, gene mapping, accession numbers and nomenclature. 6:1–42.

Ngegba PM, Khalid MZ, Jiang W, Zhong G. 2025. An overview of insecticide resistance mechanisms, challenges, and management strategies in Spodoptera frugiperda. Crop Prot. 197:107322.

Olazcuaga L, et al. 2020. A whole-genome scan for association with invasion success in the fruit fly Drosophila suzukii using contrasts of allele frequencies corrected for population structure. Mol. Biol. Evol. 37:2369–2385.

Olson ND, et al. 2023. Variant calling and benchmarking in an era of complete human genome sequences. 24:464–483.

Parmesan C, Yohe G. 2003. A globally coherent fingerprint of climate change impacts across natural systems. Nature 421:37–42.

Pedersen BS, Quinlan AR. 2018. Mosdepth: quick coverage calculation for genomes and exomes. Bioinformatics 34:867–868.

Pedersen BS, Quinlan AR. 2019. Duphold: scalable, depth-based annotation and curation of high-confidence structural variant calls. Gigascience 8(4): giz040.

Pélissié B, et al. 2022. Genome resequencing reveals rapid, repeated evolution in the Colorado potato beetle. Mol. Biol. Evol. 39:msac016.

Perini CR, et al. 2021. Transcriptome analysis of pyrethroid-resistant Chrysodeixis includens (Lepidoptera: Noctuidae) reveals Overexpression of metabolic detoxification genes. J. Econ. Entomol. 114:274–283.

Pritchard JK, Pickrell JK, Coop G. 2010. The genetics of human adaptation: hard sweeps, soft sweeps, and polygenic adaptation. Curr. Biol. 20:R208–R215.

Puechmaille SJ. 2016. The program structure does not reliably recover the correct population structure when sampling is uneven: subsampling and new estimators alleviate the problem. Mol. Ecol. Resour. 16:608–627.

Puinean AM, et al. 2010. Amplification of a cytochrome P450 gene is associated with resistance to neonicotinoid insecticides in the aphid Myzus persicae. PLoS Genet. 6:e1000999.

Pu J, Chung H. 2024. New and emerging mechanisms of insecticide resistance. Curr. Opin. Insect Sci. 63:101184.

Quero C, et al. 1997. Reinvestigation of female sex pheromone of processionary moth (Thaumetopoea pityocampa): No evidence for minor components. J. Chem. Ecol. 23:713–726.

Ranasinghe C, Headlam M, Hobbs AA. 1997. Induction of the mRNA for CYP6B2, a pyrethroid inducible cytochrome p450, inHelicoverpa armigera (Hubner) by dietary monoterpenes. Arch. Insect Biochem. Physiol. 34:99–109.

R Core Team. 2019. R: A Language and Environment for Statistical Computing. Vienna, Austria Available from: https://www.R-project.org/

Rivière J. 2011. Les chenilles processionnaires du pin: évaluation des enjeux de santé animale. Available from: https://theses.vet-alfort.fr/telecharger.php?id=1356

Robinet C, et al. 2025. Déplacement de la processionnaire du pin, Thaumetopoea pityocampa. In: Robinet C, Saintonge F-X, Tassus X, Brault S, editors. Invasion et expansion d’insectes bioagresseurs forestiers: Quels risques pour la forêt française dans le contexte des changements globauxÉditions Quae. p. 62–73.

Robinet C, Laparie M, Rousselet J. 2015. Looking beyond the large scale effects of global change: Local phenologies can result in critical heterogeneity in the pine processionary moth. Front. Physiol. 6:334.

Rosenheim JA, Johnson MW, Mau RFL, Welter SC, Tabashnik BE. 1996. Biochemical peradaptations, founder events, and the evolution of resistance in arthropods. J. Econ. Entomol. 89:263–273.

Rossi J-P, et al. 2025. Warmer and brighter winters than before: Ecological and public health challenges from the expansion of the pine processionary moth (Thaumetopoea pityocampa). Sci. Total Environ. 978:179470.

Rostant WG, et al. 2017. Pleiotropic effects of DDT resistance on male size and behaviour. Behav. Genet. 47:449–458.

Salvato P, et al. 2005. Do sexual pheromone traps provide biased information of the local gene pool in the pine processionary moth?: Pheromone traps and genetic diversity. Agric. For. Entomol. 7:127–132.

Scheffers BR, et al. 2016. The broad footprint of climate change from genes to biomes to people. Science 354:aaf7671.

Schiffels S, Wang K. 2020. MSMC and MSMC2: The Multiple Sequentially Markovian Coalescent. Methods Mol. Biol. 2090:147–166.

Scott JG. 1999. Cytochromes P450 and insecticide resistance. Insect Biochem. Mol. Biol. 29:757–777.

Secomandi S, et al. 2025. Pangenome graphs and their applications in biodiversity genomics. Nat. Genet. 57:13–26.

Shahriari M, Zibaee A, Sahebzadeh N, Shamakhi L. 2018. Effects of α-pinene, trans-anethole, and thymol as the essential oil constituents on antioxidant system and acetylcholine esterase of Ephestia kuehniella Zeller (Lepidoptera: Pyralidae). Pestic. Biochem. Physiol. 150:40–47.

Sømme L. 1989. Adaptations of terrestrial arthropods to the alpine environment. Biological Reviews 64:367–407.

Tan A, Abecasis GR, Kang HM. 2015. Unified representation of genetic variants. Bioinformatics 31:2202–2204.

Tarasov A, Vilella AJ, Cuppen E, Nijman IJ, Prins P. 2015. Sambamba: fast processing of NGS alignment formats. Bioinformatics 31:2032–2034.

Thomas CD, et al. 2004. Extinction risk from climate change. Nature 427:145–148.

Torres-Banda V, et al. 2022. Gut transcriptome of two bark beetle species stimulated with the same kairomones reveals molecular differences in detoxification pathways. Comput. Struct. Biotechnol. J. 20:3080–3095.

Tuttle AM, Hoffman TL, Schilling TF. 2014. Rabconnectin-3a regulates vesicle endocytosis and canonical Wnt signaling in zebrafish neural crest migration. PLoS Biol. 12:e1001852.

Vasseur P, et al. 2022. Human exposure to larvae of processionary moths in France: study of symptomatic cases registered by the French poison control centres between 2012 and 2019. Clin. Toxicol. 60:231–238.

Wondji CS, Hearn J, Irving H, Wondji MJ, Weedall G. 2022. RNAseq-based gene expression profiling of the Anopheles funestus pyrethroid-resistant strain FUMOZ highlights the predominant role of the duplicated CYP6P9a/b cytochrome P450s. G3 12:jkab352.

Xu L, et al. 2020. Transcriptome analysis of Spodoptera litura reveals the molecular mechanism to pyrethroids resistance. Pestic. Biochem. Physiol. 169:104649.

