## Supplementary material for "A selective sweep likely linked to xenobiotic detoxification identified along altitudinal gradients in the pine processionary moth": Suppl. Figures

### Supplementary Figures

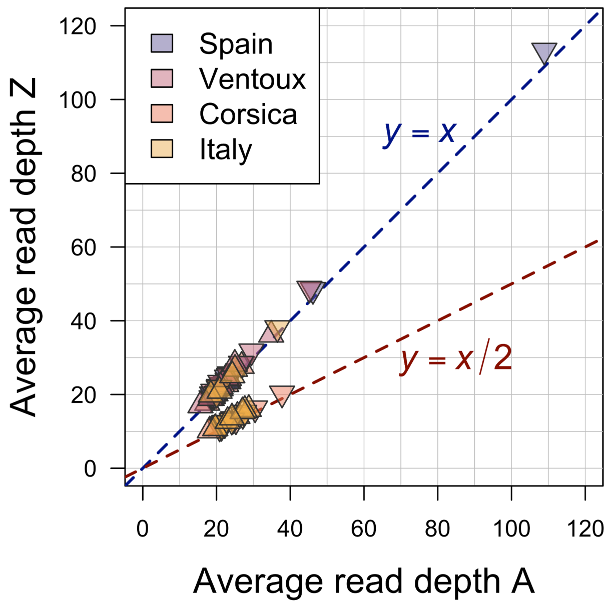


**Supplementary Figure 1.** Comparison of average sequencing depths between autosomes (x-axis) and Z chromosome (y-axis) for each ind-seq library ($n$ = 89). Triangles and inverted triangles are individuals from high and low altitude populations, respectively. In Lepidoptera, males are homogametic (ZZ), and females heterogametic (ZW). The dotted blue line gives $y = x$, as expected for male individuals ($n_{males}$ = 66). The dotted red line gives $y = x/2$, as expected for female individuals ($n_{females}$ = 23).

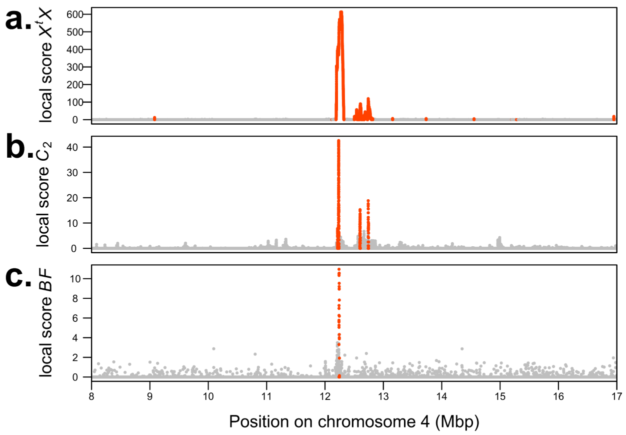


**Supplementary Figure 2.** Zoom on the major signal recovered on chromosome 4 by the different scans for signatures of selection. For each panel, SNPs located within significant local score regions are depicted in red. **a.** Local scores of the C2 statistic, contrasting low and high altitude groups of populations. **b.** Local scores of the XtX statistic measuring adaptive differentiation. **c.** Local scores of Bayes Factors (BF) measuring the strength of evidence for the linear correlation between population allele frequencies and altitude.

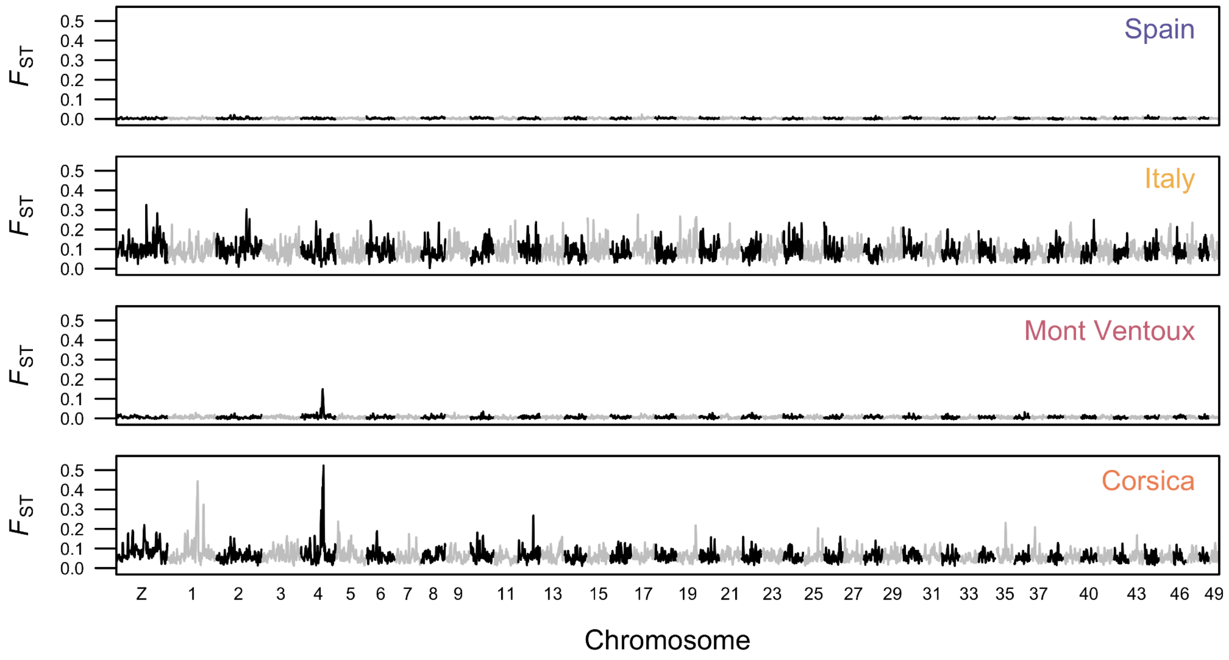


**Supplementary Figure 3.** Genome-wide pairwise $F_{ST}$ estimates computed between low and high altitude populations within each locality (given in the upper right corner of each panel) and computed from pool-seq data on 5,000 SNP windows along each chromosome (MAF ≥ 0.01).

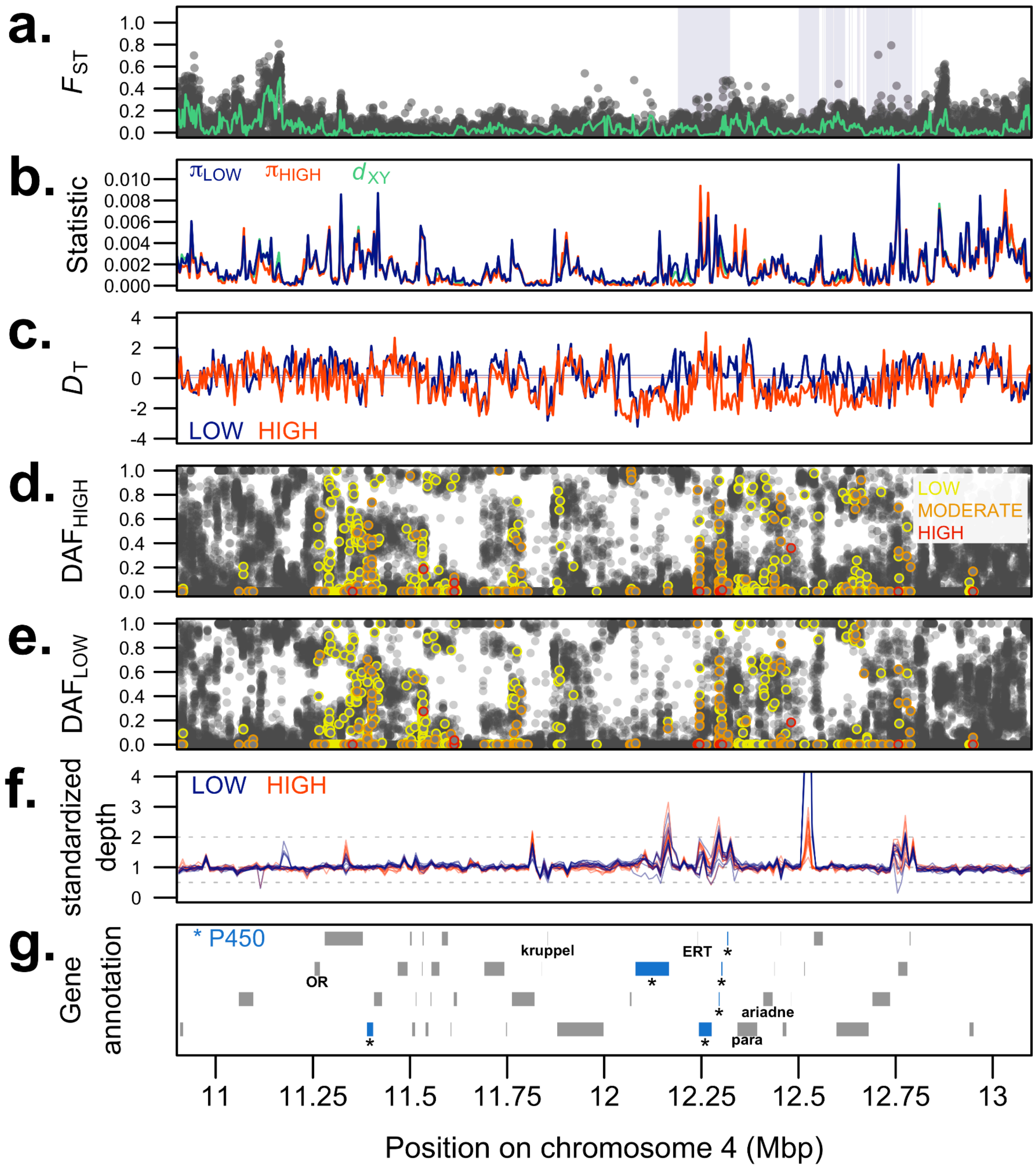


**Supplementary Figure 4.** Polymorphism and sequencing depth patterns in the vicinity of the putative selective sweep region on chromosome 4 in the Italian population pair. **a.** SNP-specific pairwise $F_{ST}$ values between low and high altitude pools (green: 100-SNP multilocus $F_{ST}$ values in sliding windows). Purple blocks indicate candidate $X^tX$ regions identified with the local score approach. **b.** Diversity within (π) and absolute divergence between ($d_{XY}$) populations, computed from ind-seq data in 5 kbp non-overlapping windows. **c.** Tajima's $D$ computed from pools in 5 kbp non-overlapping windows, with genome-wide means indicated as plain lines. **d.** Derived allele frequencies in high and **e.** low altitude pools, with SNPs colored according to their functional impact category predicted with SNPeff. **f.** Standardized sequencing depths computed from ind-seq libraries in 10 kbp windows. Dotted lines at 0.5 and 2 indicate halved and doubled average depths, respectively. **g.** Gene annotation track (cytochrome P450 genes in blue).

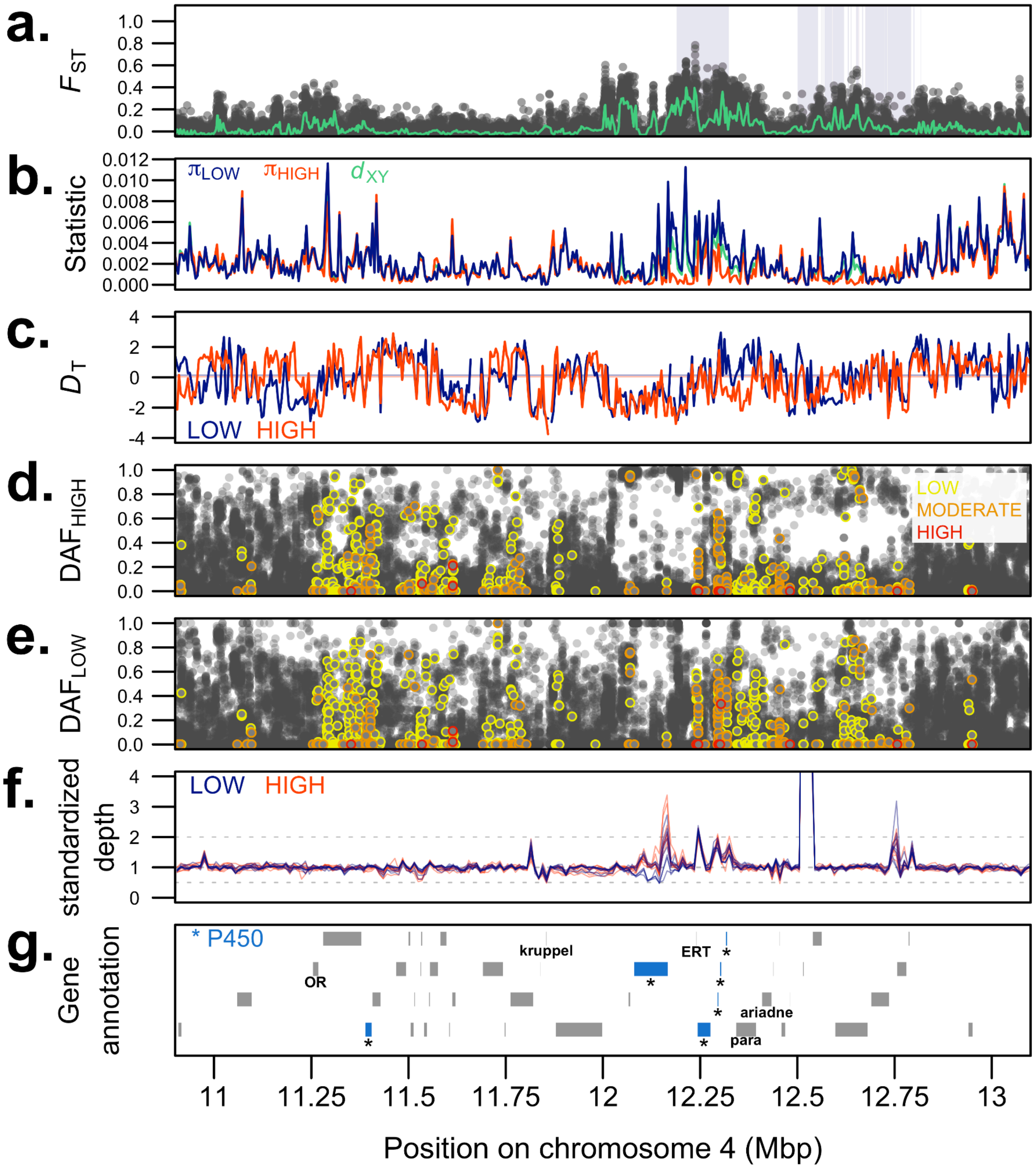


**Supplementary Figure 5.** Polymorphism and sequencing depth patterns in the vicinity of the putative selective sweep region on chromosome 4 in the Mont Ventoux population pair. **a.** SNP-specific pairwise $F_{ST}$ values between low and high altitude pools (green: 100-SNP multilocus $F_{ST}$ values in sliding windows). Purple blocks indicate candidate $X^tX$ regions identified with the local score approach. **b.** Diversity within (π) and absolute divergence between ($d_{XY}$) populations, computed from ind-seq data in 5 kbp non-overlapping windows. **c.** Tajima's *D* computed from pools in 5 kbp non-overlapping windows, with genome-wide means indicated as plain lines. **d.** Derived allele frequencies in high and **e.** low altitude pools, with SNPs colored according to their functional impact category predicted with SNPeff. **f.** Standardized sequencing depths computed from ind-seq libraries in 10 kbp windows. Dotted lines at 0.5 and 2 indicate halved and doubled average depths, respectively. **g.** Gene annotation track (cytochrome P450 genes in blue).

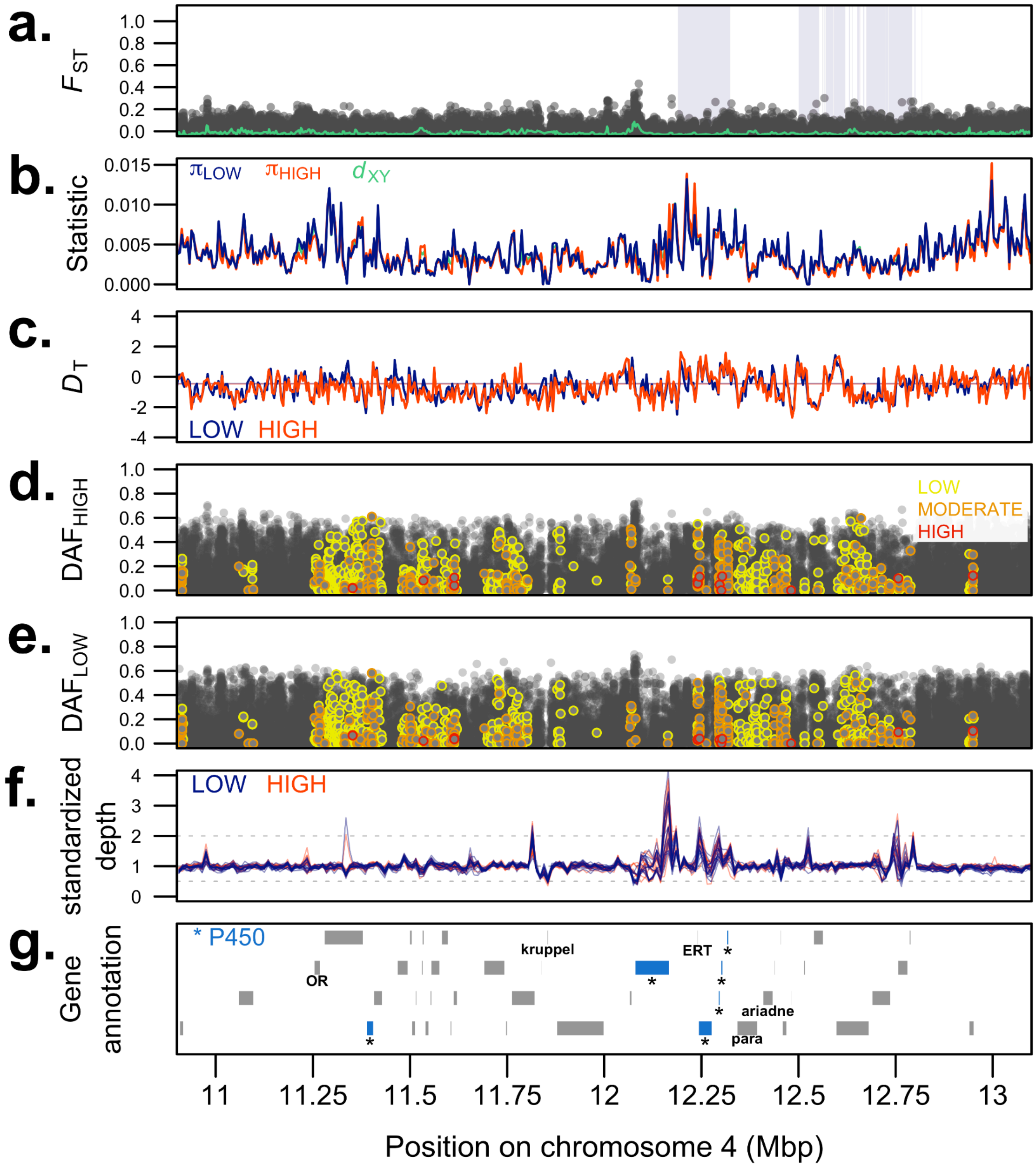


**Supplementary Figure 6.** Polymorphism and sequencing depth patterns in the vicinity of the putative selective sweep region on chromosome 4 in the Spanish population pair. **a.** SNP-specific pairwise $F_{ST}$ values between low and high altitude pools (green: 100-SNP multilocus $F_{ST}$ values in sliding windows). Purple blocks indicate candidate $X^tX$ regions identified with the local score approach. **b.** Diversity within (π) and absolute divergence between ($d_{XY}$) populations, computed from ind-seq data in 5 kbp non-overlapping windows. **c.** Tajima's *D* computed from pools in 5 kbp non-overlapping windows, with genome-wide means indicated as plain lines. **d.** Derived allele frequencies in high and **e.** low altitude pools, with SNPs colored according to their functional impact category predicted with SNPeff. **f.** Standardized sequencing depths computed from ind-seq libraries in 10 kbp windows. Dotted lines at 0.5

and 2 indicate halved and doubled average depths, respectively. **g.** Gene annotation track (cytochrome P450 genes in blue).

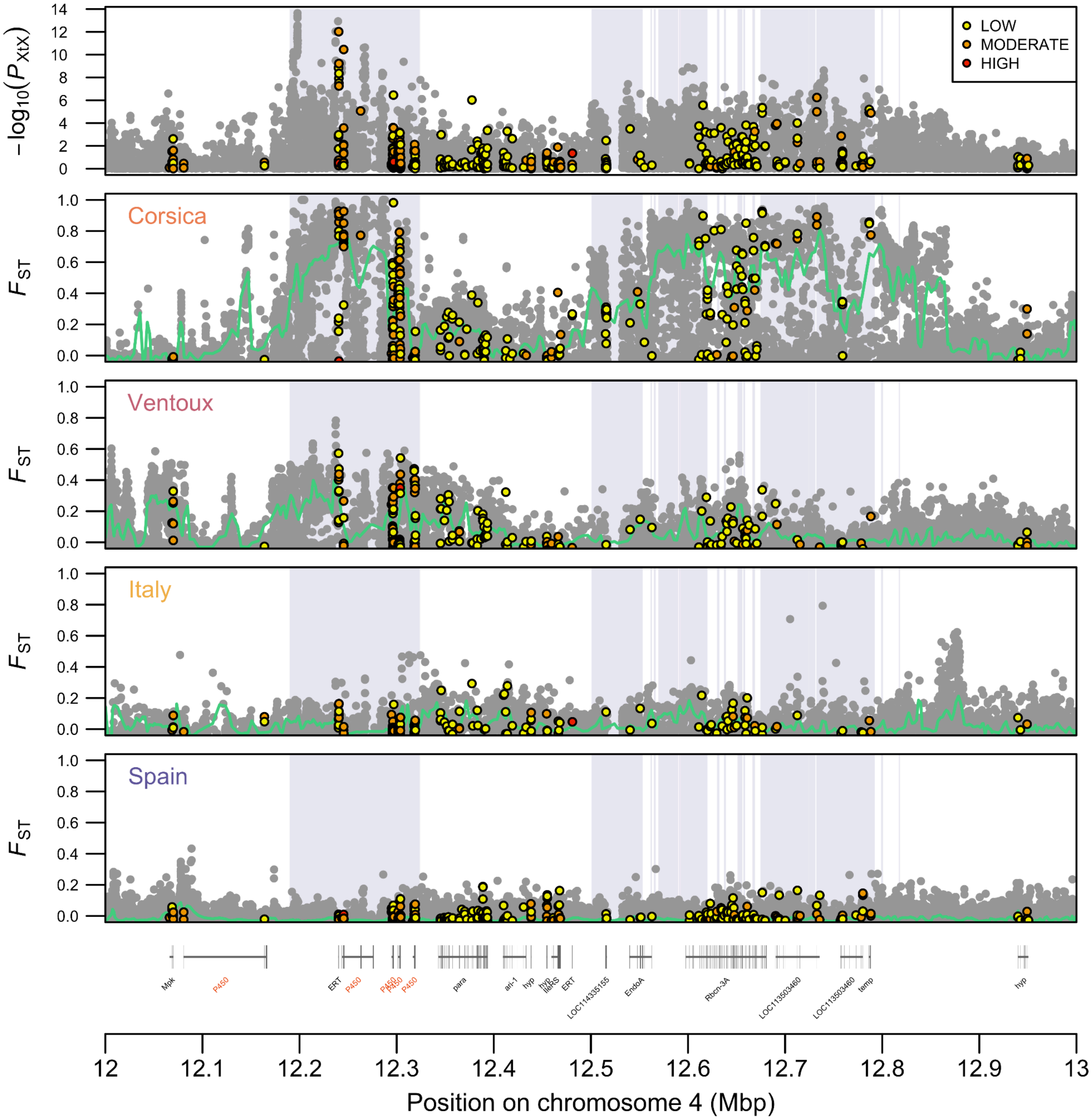


**Supplementary Figure 7.** Zoom on the gene content within the 12-13 Mb interval on chromosome 4. The upper panel gives SNP-specific $X^tX$ statistics measuring divergent selection, irrespective of the elevation levels (ξ = 3, significance level of 0.001), and the four remaining panels depict SNP-specific pairwise $F_{ST}$ values between low and high altitude pools for each locality (green: 100-SNP multilocus $F_{ST}$ values in sliding windows). Purple blocks indicate candidate $X^tX$ regions identified with the local score approach. SNPs within coding regions are colored depending on their functional impacts predicted with SNPeff. The gene annotation track is given at the bottom of the plot, with cytochrome P450 genes in red (hyp: hypothetical proteins).

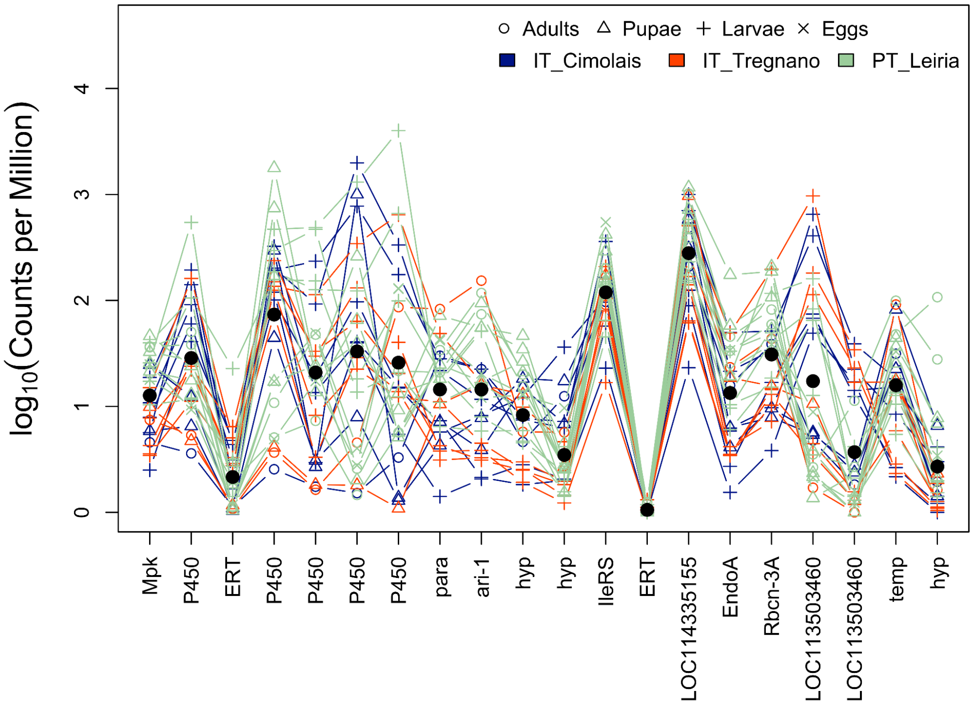


**Supplementary Figure 8.** Expression support (log-transformed counts per million reads) for the 20 genes located within the 12-13 Mb interval on chromosome 4 and ordered as in Suppl. Fig. 7. Black dots give the average over all libraries, with symbols and colors indicating developmental stages and populations, respectively (IT: Italy, PT: Portugal). See Gautier et al. (2025) and references therein for details about the RNA-seq data and its processing.

### Supplementary Table legends

**Supplementary Table 1.** Metadata and summary statistics associated with the 97 samples sequenced in the study. INSIZE: mean insert size, PERCMAP: mapping rate, DP: sequencing depth, RHO_ZA: Z-to-autosome depth ratio.

**Supplementary Table 2.** Location of the 23 genomic regions identified with the local score approach applied to $C_2$ statistics (contrast between low and high populations). LINDLEY_PEAK_VALUE: value of the highest local score peak in the region.

**Supplementary Table 3.** Gene content within the 23 genomic regions identified with the local score approach applied to $C_2$ statistics (contrast between low and high populations). Gene IDs refer to the Tpit v2.1 genome annotation and gene functions were derived from an automated Blast2GO annotation.

**Supplementary Table 4.** Location of the 745 genomic regions identified with the local score approach applied to $X^tX$ statistics (local adaptation signatures). LINDLEY_PEAK_VALUE: value of the highest local score peak in the region.

**Supplementary Table 5.** Gene content within the 745 genomic regions identified with the local score approach applied to $X^tX$ statistics (local adaptation signatures). Gene IDs refer to the Tpit v2.1 genome annotation and gene functions were derived from an automated Blast2GO annotation.

**Supplementary Table 6.** Location of the single genomic region identified with the local score approach applied to *Bayes Factors* (association with altitude). LINDLEY_PEAK_VALUE: value of the highest local score peak in the region.

**Supplementary Table 7.** Gene content within the single genomic region identified with the local score approach applied to *Bayes Factors* (association with altitude). Gene IDs refer to the Tpit v2.1 genome annotation and gene functions were derived from an automated Blast2GO annotation.

**Supplementary Table 8.** Detailed site-specific statistics for all SNPs in the 11-13 Mbp interval on chromosome 4. Columns are described in the header.
